# Implementation of CRISPR-Cas13a system in fission yeast and its repurposing for precise RNA editing

**DOI:** 10.1101/284661

**Authors:** Xinyun Jing, Bingran Xie, Longxian Chen, Niubing Zhang, Yiyi Jiang, Hang Qin, Pei Hao, Sheng Yang, Xuan Li

**Author notes:** Corresponding author (XL), (SY), (PH). These authors contributed equally to this work (Xinyun Jing, Bingran Xie).

## Abstract

In contrast to genome editing that introduces genetic changes at DNA level, disrupting or editing genes’ transcripts provides a distinctive approach to perturb a genetic system, offering benefits complementary to classic genetic approaches. To develop a new toolset for manipulation of RNA, we first implemented a member of type VI CRISPR systems, Cas13a from *Leptotrichia shahii* (LshCas13a) in *Schizosaccharomyces pombe*, an important model organism employed by biologists to study key cellular mechanisms conserved from yeast to humans. While it was shown to knock down targeted endogenous genes’ transcripts, differently from previous studies in *E. coli*, no collateral cleavage of other non-specific RNA by activated Cas13a-crRNA complex was detected in fission yeast. Second, we engineered a RNA-editing system by tethering an inactive form of LshCas13a (dCas13) to the catalytic domain of human Adenosine Deaminase Act on RNA 2 (hADAR2d), which was shown to be programmable with crRNA to target messenger RNAs and precisely edit specific nucleotide residues. We optimized the system parameters using a dual-florescence reporter and demonstrated its utility in editing of randomly selected endogenous genes’ transcripts. Our engineered RNA-editing system enables a new toolset for transcriptomic manipulation that is widely applicable in basic genetic and biotechnological research.

## Introduction

Modulation of genes’ expression or altering their transcripts provides a critical layer of regulation at RNA level in living cells. Editing or partially disrupting transcripts serves as a distinctive approach for analysis of gene functions, offering benefits complementary to classic genetic approaches, and possibly novel insight into gene function regulation at RNA level.

In recent years many molecular tools utilizing CRISPR systems to disrupt genes or introduce coding changes at DNA level were successfully developed(1-3). From a different perspective a synthetic tool to perturb a genetic system by disrupting or editing genes’ transcripts is desirable. The discovery of the type VI CRISPR systems(4-6), e.g. members of the Cas13 family, created an excellent opportunity for a new toolset for RNA-manipulation experiments. Members of the Cas13 family displayed a unique ability of targeting single-strand RNA (ssRNA). They cleave RNA at sites guided by a CRISPR RNA (crRNA) containing a variable length spacer for different enzyme(7-10). A member of Cas13 family, Cas13a from *Leptotrichia shahii* (LshCas13), was found to knockdown target transcripts in *E. coli* and mammalian cells, and was used in nucleic acid detection and RNA tagging(7,8,11,12). More recently, a different member of Cas13 family, Cas13b from *Prevotella sp. P5-125*, was used to construct a fusion protein for targeted editing of genes’ transcripts in mammalian cells, which was able to correct disease-relevant mutations in human cell lines(13).

The unicellular organism, *Schizosaccharomyces pombe* (fission yeast) is one of the popular and important model systems employed by biologists to study processes that are conserved from yeast to humans. Its usage in research led to many key discoveries in cell cycle control(14,15), chromosome structure(16,17), histone modifications (18,19), cytokinesis(20,21), etc. Along the way, genetic manipulation tools were developed and applied, having an exceptional impact on the scientific community. To take advantage of the new advance in CRISPR research, our current study was set out to explore a synthetic tool set for manipulation of gene transcripts in fission yeast by implementation and repurposing of the type VI CRISPR system.

There are unique advantages associated with manipulation of transcripts in stead of genes. Altering genes’ transcripts does not make any change to genes themselves, making it reversible and more temporally and spatially controllable. It is also more efficient and effective to operate for genes of multiple copies, especially in polyploid organisms. It is a potential useful alternative to disruption or correction of mutated genes in diseased tissues for therapeutic purpose. In addition, implementation of a RNA manipulating system in fission yeast has added benefits for studying cellular functions/ processes because of its close similarity to higher eukaryotic cells. There is no native protein like Adenosine Deaminase Act on RNA (ADAR) family of enzymes(22), in fission yeast to interfere with engineered RNA editing system. Partially editing a gene’s transcripts allows one to simulate the effect of two different alleles for a given gene in haploid yeast strains. For genetic screening experiments in haploid fission yeast, a conditional knockdown or editing of genes’ transcripts may circumvent the problem of lethal effect for some mutations, which is easier to achieve than using traditional genetic methods.

Motivated by these purposes, in the current study we first sought to implement the LshCas13a system in *S. pombe* for targeting gene transcripts, before we could use it to engineer a programmable system for editing genes’ transcripts at specific sites. Similar to the mutant Cas9 variant (dCas9) capable of binding target DNA but inactive in DNA cleavage(23,24), one of the four LshCas13a HEPN-domain mutants, R1278A mutant (termed dCas13a), had stronger target RNA binding (K_D_ ∼ 7 nM) than the wild type (K_D_ ∼ 46 nM), albeit being catalytically inactive(7). Note the binding affinity on target RNA by the R1278A mutant complex in presence of crRNA was one magnitude higher than that of crRNA binding to target RNA (K_D_ ∼ 69 nM)(7). So we designed and engineered an RNA-editing system by tethering the inactive form of LshCas13a (R1278A mutant) (dCas13a), to the catalytic domain of human Adenosine Deaminase Act on RNA 2 (hADAR2d). We then evaluated the utility of dCas13a in targeting specific mRNA for precise RNA editing in fission yeast. We showed *in vivo* that this fusion complex can be programmed to target genes’ transcripts and precisely edit specific nucleotide residues in presence of crRNA. We optimized the system parameters and demonstrated its utility in editing of randomly selected endogenous gene transcripts in addition to constructed fluorescent reporter transcripts. Our work enables a new programmable toolset in fission yeast for transcriptomic manipulation that is widely applicable in basic genetic and biotechnological research.

## Materials and Methods

### Plasmids and constructs

Plasmids and constructs are listed in Tables S1, S2, S4, S5 and S6. Note those obtained from the Addgene repository (http://www.addgene.org) are indicated. All sequences of oligonucleotides used in the study are described in Dataset S1. For molecular cloning, *E*.*coli* DH5α was used.

PCR was performed using Taq (Thermo Scientific) or KOD FX DNA polymerase (TOYOBO). Plasmids and chromosomal DNA were extracted using Plasmid Mini Kit I and Gel Extraction Kit from OMEGA. Cloning was performed using restriction endonucleases and T4 DNA ligase (New England Biolabs), or ClonExpress ® II One Step Cloning Kit (Vazyme).

Plasmids pDUAL-HFF1-Cas13a, pDUAL-HFF1-dCas13a and pDUAL-HFF1-Cas13a-hADAR2d were generated by inserting the PCR fragments of LshCas13a, dCas13a, and dCas13a-hADAR2d into the NdeI/NcoI restriction sites of pDUAL-HFF1. The Cas13a gene fragment was amplified using plasmid pC001 (Table S1) as template and primers Cas13a-P5 /Cas13a-P3. The dCas13a gene fragment was obtained by overlap PCR using plasmid pC001 as template and primers Cas13a-P5, Cas13a-mut-mid-P3, Cas13a-mut-mid-P5 and Cas13a-P3. The dCas13a-hADAR2d gene fragment was obtained by overlap PCR using dCas13a gene fragment and ADAR2 gene fragment as template and primers Cas13a-P5, XTEN-dCas13a-P3, XTEN-hADAR2d-P5, hADAR2d-P3.

To generate plasmids pDUAL-HFF1-eGFP, pDUAL-HFF1-mCherry-eGFP and pDUAL-HFF1-mCherry-eGFP-W58X, fragments of eGFP, mCherry-eGFP and mCherry-eGFP-W58X were obtained with from PCR amplification or digestion of intermediate cloning plasmids pKS-mCherry-eGFP or pKS-mCherry-eGFP-W58X (Table S2), and were cloned into the NheI/BglII sites in plasmid pDUAL-HFF1. To produce the intermediate cloning plasmids pKS-mCherry-eGFP and pKS-mCherry-eGFP-W58X, we first amplified the mCherry-linker fragment by overlap PCR using plasmid pmCherry Paxillin (Addgene: 50526) and plamid pLinker (synthesized by Genwiz, Suzhou, China) as templates using primers mCherry-P5, mCherry-linker-P3, mCherry-linker-P5 and linker-P3. The mCherry-linker gene fragment was cloned into the KpnI/XbaI sites of plasmid pBluescript II KS(+) (simplified as pKS in Table S1) producing intermediate plasmid pKS-mCherry-linker. Second, we amplified eGFP gene fragment using plasmid pEGFP-N1-FLAG (Addgene: 60360) as template with primers eGFP-P5/ eGFP-P3. Meanwhile we amplified the eGFP-W58X gene fragment by overlap PCR using plasmid pEGFP-N1-FLAG as template with primers eGFP-P5, eEGFP-mut-P3, eEGFP-mut-P5, eGFP-P3. Third, the eGFP and eGFP-W58X gene fragment were cloned into XbaI/SacI sites of plasmid pKS-mCherry-linker generating plasmids of pKS-mCherry-eGFP and pKS-mCherry-eGFP-W58X.

The intermediated crRNA cloning construct was built by synthesizing the gene composing *rrk1* promoter, leader RNA, BspQI placeholder, HDVR Ribozyme, BsaI placeholder and Hammerhead Ribozyme(synthesized by Genwiz, Suzhou, China), and digesting the synthesized DNA with ClaI/EcoRV restriction enzyme, before cloning them into the pKS plasmid yielding pKS-crRNA-backbone vector. Complementary primers used for generation of crRNA and pRNA were combined (5 μl of 100 μM solution) in Tris buffer and annealed by heating to 95 °C for 5 min, followed by a gradual cooling to 45 °C at a rate of 0.1 °C per second to generate dsDNA substrates with sticky end. crRNA were cloned into BspQI placeholder of plasmid pKS-gRNA-backbone, producing intermediate plasmids in Table S2. pRNA was cloned into the *Bsa*I placeholder of plasmid pKS-rrk1-crRNA-control yielding plasmid pKS-rrk1-crRNA-pRNA-separate as Table S2.

The crRNA expressing plasmids was constructed using ClonExpress ® II One Step Cloning Kit. The crRNA fragments were amplified using the intermediated crRNA cloning construct (as in Table S2) as templates using primers pDUAL-SpeI-T3 and pDUAL-XhoI-T7. The plasmids pDUAL-HFF-mCherry-eGFP-W58X, pDUAL-HFF1-Cas13a, and pDUAL-HFF1 were digested with SpeI and PspXI. The pDUAL-HFF1-Cas13a SpeI/PspXI fragment and the crRNA-ade6-1160, crRNA-tdh1-79 PCR fragments were recombined using ClonExpress ® II One Step Cloning Kit generating plasmids as in Table S4. pDUAL-HFF-mCherry-eGFP-W58X SpeI/PspXI fragment and crRNA-28bp, crRNA-37bp, crRNA-39BP, crRNA-49bp, crRNA-59bp, crRNA-65bp, crRNA-(−9)-65, crRNA-(−5)-65, crRNA-(−4)-65, crRNA-2-65, crRNA-12-65, crRNA-15-65, crRNA-19-65 PCR fragments were recombined using ClonExpress ® II One Step Cloning Kit, generating plasmids as in Table S5. pDUAL-HFF-mCherry-eGFP-W58X SpeI/PspXI fragment and crRNA-act1-1566, crRNA-ade6-622, crRNA-ade6-1002, crRNA-ade6-1160, crRNA-erp5-672, crRNA-mel1-921, crRNA-mug45, crRNA-nmt1-648, crRNA-tdh1-79 PCR fragments were recombined using ClonExpress ® II One Step Cloning Kit generating plasmids as in Table S6.

### Strains and transformation

The *S. pombe* strain FY7652 containing the *ura4*-D18 and *leu1*-32 alleles was used in this study (Table S1). Strains were grown in YES medium(25) supplemented with 50 μg/ml uracil and 50μg /ml leucine at 30 °C to mid-log phase and transformed using the Lithium Acetate/PEG/Heat shock method(25). For chromosomal expression of eGFP or dCas13a-hADAR2d, plasmids pDUAL-HFF-eGFP or pDULA-HFF1-dCas13a-hADAR2d were digested with NotI restriction enzyme and treated with FastAP thermosensitive alkaline phosphatase (Thermo Scientific), before DNA was recovered with Gel Extraction Kit (OMEGA). Transformation was carried out with 500 ng recovered linear DNA, i.e. pDUAL-HFF-mCherry-eGFP or pDULA-HFF1-dCas13a-hADAR2d. Transformants were selected on EMM medium supplemented with 50 μg/ml uracil. For episomal expression of Cas13a constructs (alone or together with crRNA), mCherry-eGFP-W58X constructs (alone or together with crRNA), or mCherry-eGFP construct (as positive control), 100 ng circular plasmid was used to transformed various yeast strains and selected on EMM medium without supplements.

### Measuring growth rate for different *S. pombe* strains

The *S. pombe* strain, FY7652, transformed with plasmids pDUAL-HFF1, pDUAL-HFF1-Cas13a, pDUAL-HFF1-Cas13a-tdh1, or pDUAL-HFF1-Cas13a-ade6 were plated in EMM(25), and grew for 4 days. Colonies were picked and used to seed cultures of 3 mL EMM medium, growing until mid-log phase. Harvested cells were inoculated in 20 mL EMM medium with starting optical density (OD600) at 0.1. OD600 was measured for each culture for different time points during cell growth.

For comparison of growth rate on plates, yeast cells (FY7652) carrying empty plasmid (PDUAL-HFF1) or plasmid encoding LshCas13a (pDUAL-HFF1-Cas13a) were cultured overnight. The cell cultures were then adjusted to 1 OD/ml with EMM medium. The cell suspensions were diluted at 10^−1^, 10^−2^, and 10^−3^, respectively, and 3μ l for each dilution were plated out on EMM or EMM+thiamine plates, respectively. Plated cells were allowed to grow for two days before being photographed.

### Measuring transcript levels by quantitative PCR

To measure transcript levels in *S. pombe* cells, strains carrying plasmids pDUAL-HFF1-Cas13a, pDUAL-HFF1-Cas13a-tdh1, or pDUAL-HFF1-Cas13a-ade6 were cultured to late log phase. Yeast cells were harvested before total RNA was extracted using RNeasy Plus Mini Kit (Qiagen) and digested with DNase I to remove possible contamination by genome DNA. Reverse transcription was performed on total RNA using the RevertAid First Strand cDNA Synthesis Kit (Thermo Scientific) and oligodT primers. Gene transcript levels were quantified with qPCR using TaKaRa_SYBR® *Premix Ex Taq*™ II_RR820Q kit (TAKARA) following the manufacture’s instructions. The primer sets, tdh1-79-RT-p5/tdh1-79-RT-p3, ade6-1160-RT-p5/ade6-1160-RT-p3, and Act1-1566-RT-p5/Act1-1566-RT-p3 were used for amplification of *tdh1, ade6*, and *act1*, respectively (Dataset 1). Note *act1* was used as a reference for determining the relative transcript level of *tdh1* and *ade6*. All qPCR reactions were performed in 20-μl reactions with three technical replicates in 96-well plates, using StepOnePlus™ Real-Time PCR System (Thermo Fisher Scientific). To normalize RNA inputs, ΔCt values for each sample (e.g. culture of strain carrying plasmid pDUAL-HFF1-Cas13a) were computed by subtracting Ct values for *act1* from Ct values for targets (e.g. *tdh1*). The relative transcript abundance (normalized against *act1*) for each target was obtained as 2^-^ΔCt. The relative transcript abundance from control sample (strain carrying plasmid pDUAL-HFF1-Cas13a) was set as standard ‘1’ (Fig. 1*D*); and the normalized expression of *tdh1* for strain carrying plasmid pDUAL-HFF1-Cas13a-rrk1-crRNA-tdh1, or of *ade6* for strain carrying plasmid pDUAL-HFF1-Cas13a-rrk1-crRNA-ade6 were calculated by comparing against control samples.

**Figure 1.**
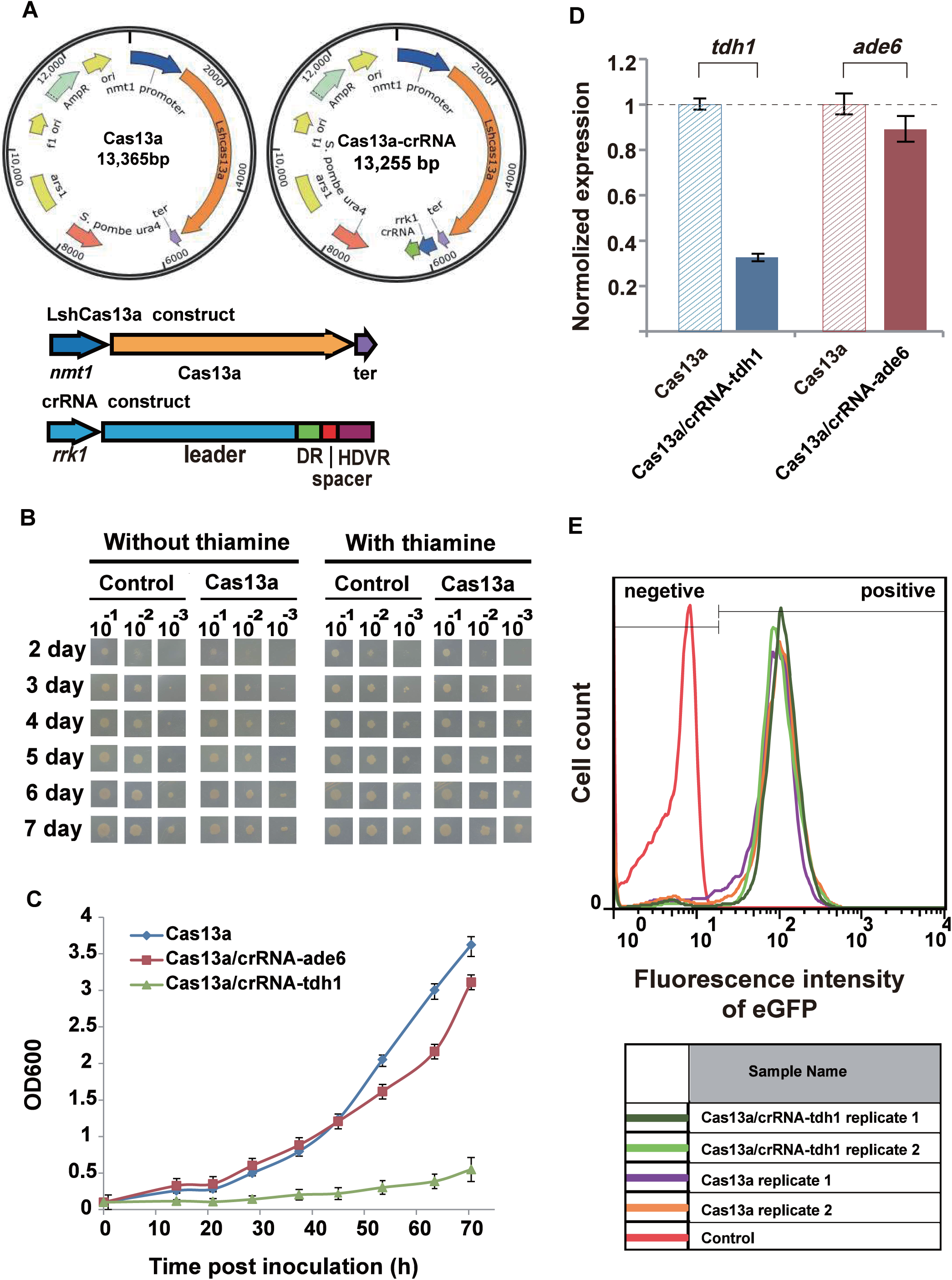
Implementation of LshCas13a system in fission yeast. (A) An *E. coli*-*S. pombe* shuttle plasmid (pDUAL-HFF1(28)) was modified to contain either LshCas13a construct or both LshCas13a and crRNA constructs in the same plasmid. The LshCas13a sequence is driven by *nmt1* promoter, and crRNA by *rrk1* promoter. The *rrk1* transcription unit was combined with HDVR to generate clean crRNA(27). DR, direct repeat region of crRNA; HDVR, Hepatitis Delta Virus Ribozyme; ter, *ADH1* terminator. (B) Growth phenotype of *S. pombe* strain, FY7652, transformed with empty plasmid or plasmid encoding LshCas13a only. Cells carrying empty plasmid or plasmid encoding LshCas13a were cultured overnight and adjusted to 1 OD/ml with EMM medium. Cell suspensions were diluted at 10^−1^, 10^−2^, and 10^−3^, and 3μl for each dilution were plated out. Control, empty plasmid (pDUAL-HFF1); Cas13a, plasmid encoding LshCas13a. (C) Growth curve of *S. pombe* strains expressing LshCas13a alone, or LshCas13a together with designed crRNA targeting *tdh1* (crRNA-tdh1) or *ade6* (crRNA-ade6) transcripts. All values are mean ± s.e.m. with n=2. (D) Knockdown effect on *tdh1* or *ade6* genes’ transcripts by LshCas13a in *S. pombe*. qPCR was performed for samples Cas13a (strain carrying plasmids pDUAL-HFF1-Cas13a), Cas13a/crRNA-tdh1(carrying pDUAL-HFF1-Cas13a-rrk1-crRNA-tdh1), and Cas13a/crRNA-ade6 (carrying pDUAL-HFF1-Cas13a-rrk1-crRNA-ade6). Normalized expression values of target genes for each sample were deduced as described (*Materials and Methods*). All values are mean ± s.e.m. with n = 4 unless noted otherwise. (E) Distribution of florescence intensity in *S. pombe* cells. Control, *S. pombe* (FY7652); Cas13a, *S. pombe* strain (with chromosomal eGFP gene) transformed with pDUAL-HFF1-Cas13a; Cas13a/crRNA-tdh1, *S. pombe* strain (with chromosomal eGFP gene) transformed with pDUAL-HFF1-Cas13a-rrk1-crRNA-tdh1.

### Detection of expression levels for endogenous eGFP in *S. pombe* using flowcytometry

To measure green fluorescence intensity of *S. pombe* cells, 20 ml cultures of wild-type *S. pombe* strain (FY7652), and strains (with chromosomal eGFP gene) carrying plasmids pDUAL-HFF1-Cas13a, pDUAL-HFF1-Cas13a-rrk1-crRNA-tdh1, and pDUAL-HFF1-Cas13a-rrk1-crRNA-ade6, were grown to late log phase. Cells were harvested, and approximately 1×10^7^ cells were fixed with 4% formaldehyde for two hours. Cells were then washed for five times with PBS buffer and resuspended in 1 mL PBS buffer. Florescence intensities for cells of different strains were measured with BD LSR II SORP flow cytometer (BD Biosciences), with configuration: 488 nm blue laser, 505 LP filter, and 530/30 BP filter with PMT position on E. Raw data were analyzed with Flowjo7.6 software (Becton, Dickinson & Company).

### Visualization of red (mCHERRY) and green (eGFP) florescence in *S. pombe* cells

*S. pombe* strains expressing dCas13a-hADAR2d, and transfected with different plasmids were grown in EMM medium at 30 °C to exponential growth phase. Cells were harvested by centrifugation for 5 min at 1200 g at room temperature, washed with 500 ul of distilled water twice and resuspended in distilled 60% glycerol at 0.5-1.0×10^7^ cells/100 ul. 2.5 ul cell suspensions were placed on a 60×24 mm cover slip coated with poly-L-Lysine and covered with 18×18 mm cover slip. Fluorescence image was visualized using ZEISS LSM 880 Confocal Laser Scanning Microscope (Axio imager2) with C-Apochromat 40×/1.2 W KorrM27 objective and PMT detector. The excitation and emission wavelength for red florescence (mCherry) is 543 and 621 nm, and that for green florescence (eGFP) is 488 and 523 nm.

### Estimating RNA editing efficiency

To obtain florescence intensities of yeast cells, 20 ml cultures of *S. pombe* strains (with chromosomal dCas13a-hADAR2d gene) carrying different plasmid constructs, i.e. positive control (mCherry-eGFP), negative control (mCherry-eGFP-W58X), or mCherry-eGFP-W58X with various crRNA/pRNA configurations (Fig. 2*B*), were grown to late log phase. Cells were harvested, and approximately 1×10^7^ cells were fixed with 4% formaldehyde for two hours. Cells were then washed for five times with PBS buffer and resuspended in 1 mL PBS buffer. Florescence intensities at two channels, i.e. 488 and 355 nm, were measured with the BD LSR II SORP flow cytometer (BD Biosciences) with configuration: 488 nm blue laser, 505LP LP filter, and 530/30 BP filter with PMT position on E (for eGFP detection); and 355 nm UV laser, 600 LP filter, and 610/20 BP filter with PMT position on A (for mCHERRY detection). Raw data were analyzed with Flowjo7.5 software (Becton, Dickinson & Company).

**Figure 2.**
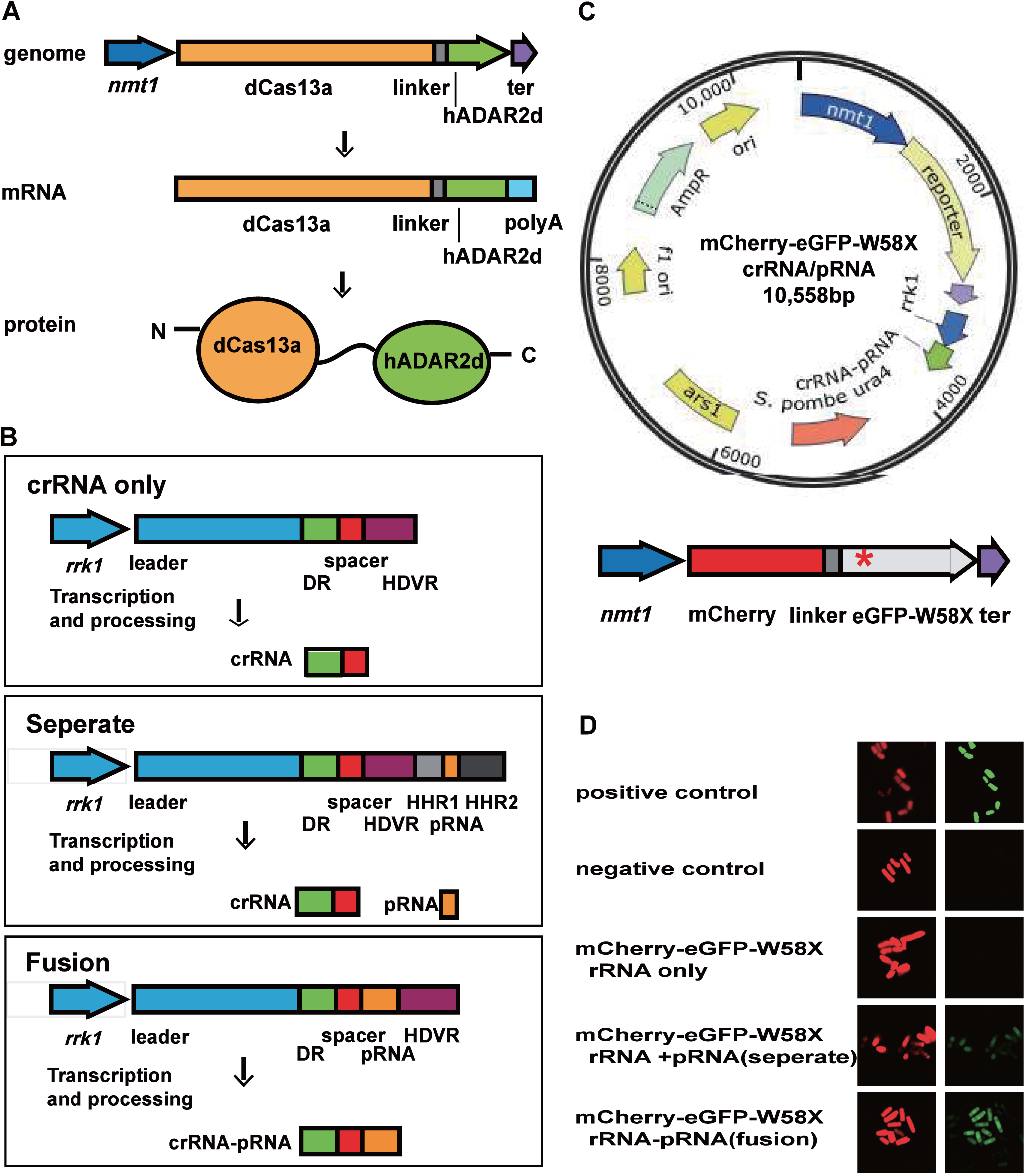
Construct and validation of dCas13a-medicated system for site-specific editing of mRNA for fluorescent reporter. (A) Scheme for chromosomal expression of dCas13a-hADAR2d in *S. pombe*. dCas13a-hADAR2d fusion gene (under *nmt1* promoter) was integrated into the *leu1* locus in *S. pombe* chromosome (*Materials and Methods*). ter, ADH1 terminator. (B) Scheme of different crRNA/pRNA expressing constructs in plasmid pDUAL-HFF1 (inserted between PspXI and SpeI). The *rrk1* transcription unit was combined with HDVR, HHR1(46), and HHR2(27) to generate clean crRNA and pRNA fragments, or their fusion molecules as illustrated. Note a mismatched nucleotide opposite to the target adenosine was introduced in pRNA(35). DR, direct repeat region of crRNA; HDVR, Hepatitis Delta Virus Ribozyme; HHR1, Hammerhead Ribozyme1; HHR2, Hammerhead Ribozyme2. (C) Scheme of fluorescent reporter construct. The dual-fluorescence reporter (mCherry and eGFP fusion gene) had a stop codon (UAG) at amino acid position 58 of eGFP (named eGFP-W58X), represented by red star (*). The reporter construct is placed in the same plasmid of the crRNA/pRNA construct as in 2b. ter, ADH1 terminator. (D) Validation of dCas13a-medicated site-specific RNA editing of the dual-fluorescent reporter. Confocal images of dCas13a-hADAR2d expressing *S. pombe* cells transfected with different plasmid constructs are visualized using ZEISS LSM 880 Confocal Laser Scanning Microscope (*Materials and Methods*). Pictures were taken from yeast cells of exponential growth phase. Red and green images are shown for the same field of cells. Positive control, mCherry-eGFP in pDUAL-HFF1; negative control, mCherry-eGFP-W58X in pDUAL-HFF1.

To estimate RNA editing efficiency by ratio of florescence intensities between eGFP and mCHERRY, i.e. the ratio of green fluorescence intensity to red fluorescence intensity, the ratio of negative control cells and positive control cells were used as baseline (0%) and maximum (100%), respectively, to normalize and calculate RNA editing efficiency for samples of different constructs (between 0-100%)

### Estimating RNA editing efficiency by Sanger sequencing

To determine editing efficiency basing on ratio of edited base (I) to unedited base (A), total RNA was extracted using RNeasy Plus Mini Kit (Qiagen) and was digested with *DNase* I to eliminate possible contamination by genome DNA. cDNA was synthesized using the RevertAid First Strand cDNA Synthesis Kit (Thermo Scientific) with oligodT primers. DNA fragments for target genes were amplified by PCR using specific primers (Dataset 1) before being sent for Sanger sequencing (GeneWiz, Suzhou). RNA editing efficiency for each editing site was estimated with the ratio of signal peak height for edited base to that for sum of edited and unedited bases, according to the published study(26).

## Results

### Implementation of the type VI CRISPR-Cas13a system in fission yeast

To implement the type VI CRISPR-Cas13a system in fission yeast, *E. coli*-*S. pombe* shuttle plasmids were constructed to carry either LshCas13a coding sequence alone (under *nmt1* promoter), or together with pre-crRNA sequences (under a separate *rrk1* promoter) on same plasmids (Fig. 1*A*). The *rrk1* promoter was previously used to generate crRNA for *Streptococcus pyogenes* Cas9 (SpyCas9) in fission yeast(27). Similarly, we included the HDV Ribozyme(27) sequence downstream of the pre-crRNA sequences, which helped remove the 3’-trailing sequences to form clean crRNA. We designed different crRNA constructs to target two endogenous genes, *tdh1* and *ade6*, in *S. pombe*, respectively.

The constructed plasmids were transformed into a *S. pombe* strain expressing endogenous eGFP (*Materials and Methods*). The transformed yeast cells were first plated out in media containing thiamine that inhibited the expression of LshCas13a under *nmt1* promoter. The strains transformed with empty plasmid (pDUAL-HFF1)(28), or the plasmid containing only LshCas13a construct were viable and grew at similar rates in presence of thiamine (Fig. 1*B*). However, when thiamine was withdrawn from media, we observed a slightly slower growth for *S. pombe* strains carrying plasmid containing LshCas13a construct, compared to those with empty plasmid (Fig. 1*B*). Slower growth was often observed when foreign proteins were expressed in yeast(29,30), which suggested possibly mild toxicity for LshCas13a effector toward *S. pombe in vivo*.

We next investigated the knockdown effect of LshCas13a with introduction of crRNAs targeting the two endogenous genes’ transcripts (Tables S3 and S4). While the presence of crRNA targeting *tdh1* transcript (crRNA-tdh1) caused significant slow growth of the yeast cells comparing to those with LshCas13a only, the crRNA targeting *ade6* (crRNA-ade6) showed no significant effect on their growth rate (Fig. 1*C*). We further determined the knockdown effect on the *tdh1* and *ade6* transcripts by performing qPCR analysis on total RNA isolated from the transformed yeast strains. When comparing to the controls expressing only LshCas13a (without crRNA), we observed a dramatic reduction in transcript level of *tdh1* gene, but less significant reduction in that of *ade6* gene (Fig. 1*D*).

Our implemented CRISPR-Cas13a system in fission yeast appeared to target genes’ transcripts with different efficiency. The observed impaired growth rates in yeast cells with reduced *tdh1* transcript, could be due to “collateral effect” of Cas13a-crRNA complex activated by binding of target RNAs, which was found by previous studies(7,8). So to investigate the possible “collateral effect” of activated LshCas13a complex in *S. pombe* cells, we looked up the expression levels of endogenous eGFP in control cells with LshCas13a only and in those with both LshCas13a and crRNA. Compared to strains expressing LshCas13a only, introduction of crRNA targeting either *tdh1*gene or *ade6* gene transcripts did not result in observed reduction of endogenous eGFP expression, evidenced by their unchanged cellular florescence intensity observed with flowcytometer (Fig. 1*E* and Fig. S3). So the non-specific RNase activity for activated Cas13a-crRNA complex was not detected in fission yeast by our experiment. The implementation of LshCas13a in *S. pombe* laid the foundation for developing a new toolset in manipulating gene transcripts in the popular model organism.

### Engineering a site-specific RNA editing system using LshCas13a

The R1278A mutant (dCas13a) of LshCas13a was found previously to lose target RNA cleavage activity but retain sequence-specific RNA binding capability(7). So we explored the possibility to utilize dCas13a to anchor a RNA deaminase domain for precise RNA editing function in *S. pombe*. As a proof of concept, we designed and engineered a fusion construct by tethering the deaminase domain of human ADAR2(31,32) (termed hADAR2d) to the C-terminus of dCas13a with a 16-amino acid linker(33) (Fig. 2*A*). Although human ADAR2 was known to catalyze hydrolytic deamination of adenosine (A) to form inosine (I) on double stranded regions of RNA substrates, its C-terminal domain (hADAR2d) had little activity toward RNA substrate without N-terminus dsRNA-binding motif(34). To test the function of dCas13a and hADAR2d fusion construct (termed dCas13a-hADAR2d) *in vivo*, we first generated the *S. pombe* strain that expressed the fusion protein, in which a single-copy of dCas13a-hADAR2d fusion gene under *nmt1* promoter was integrated into the *leu1* locus in *S. pombe* chromosome (Fig. 2*A*, and *Materials and Methods*). To guide dCas13a-hADAR2d protein to target transcripts for RNA editing, we constructed new plasmids from pDUAL-HFF1, to contain crRNAs for dCas13a targeting, and pairing RNAs (pRNA) for forming dsRNA substrate required by hADAR2d. While the crRNA and pRNA transcription cassette was driven with a *rrk1* promoter(27), three different constructs were made: 1) crRNA only as control; 2) crRNA and pRNA as two separate molecules; and 3) crRNA and pRNA as one fusion molecule (Fig. 2*B*). These plasmid constructs would remain episomal after they were transfected into the dCas13a-hADAR2d expressing *S. pombe* strain (*Materials and Methods*).

### dCas13a mediated site-specific editing of mRNA for fluorescent reporter

Our RNA editing system in fission yeast (strain FY7652) consisted of an integrated dCas13a-hADAR2d fusion gene in genome and a modified episomal vector containing crRNA/pRNA constructs. In order to facilitate testing RNA editing activity on target RNA substrates, we next constructed a reporter of dual-fluorescence fusion protein consisting of mCherry and eGFP, as described(35) but with some modifications (*Materials and Methods*). A single nucleotide mutation was introduced to generate a stop codon (UAG) at amino acid position 58 of eGFP (named eGFP-W58X) (Fig. 2*C*). Without RNA editing to correct the mutated sequence, the reporter protein would emit red fluorescence only. However, once the reporter’s eGFP-W58X mutation is repaired by our RNA editing system, it would allow translation to continue to its completion, which emits both red and green fluorescence. The dual-fluorescence fusion protein reporter made it easy to estimate the efficiency of RNA editing by quantifying and comparing fluorescence intensities at two different channels. It avoided the variation in fluorescence intensity among yeast cells with single-fluorescence protein reporter. To make it simple, we placed the reporter constructs (mCherry-eGFP-W58X driven by *nmt1* promoter) in the same plasmids containing different crRNA/pRNA constructs (driven by *rrk1* promoter) (Fig. 2*C*).

We found while dCas13a-hADAR2d expressing cells carrying negative control plasmid (mCherry-eGFP-W58X in pDUAL-HFF1) had red fluorescence only, those carrying positive control (mCherry-eGFP fusion in pDUAL-HFF1) displayed both red and green fluorescence (Fig. 2*D*, first and second rows). When only crRNA was added, still red fluorescence was detected only, indicating no RNA editing took place on reporter RNA (Fig. 2*D*, third row). However, when both crRNA and pRNA were added, both red and green fluorescence were observed for the same cells (Fig. 2*D*, fourth and fifth rows).

The results indicated that the dCas13a-hADAR2d/crRNA+pRNA system was capable of editing reporter mRNA at specific nucleic acid, which required both crRNA and pRNA for the activity. Furthermore, between the two different configurations that crRNA and pRNA were generated either as two separate RNA molecules or as one fusion molecule, the fusion construct gave an apparently stronger green fluorescent signal than that with separate crRNA and pRNA molecules, suggesting the fusion molecule had a better efficiency in either guiding dCas13a-hADAR2d to target RNA, supporting catalytic reaction of hADAR2d on dsRNA substrate, or both. The *in vivo* RNA editing efficiency of the dCas13a-hADAR2d system was measured by intensity ratio of green to red fluorescence (normalized with both positive and negative controls), and by direct sequencing of reverse-transcribed reporter transcripts. The RNA editing efficiency for crRNA-pRNA fusion construct, and for separate crRNA and pRNA molecules was estimated to be ∼7.2% and ∼0.7%, respectively (*Materials and Methods*, Dataset S1, and Fig. S1).

### Working parameter window for dCas13a mediated RNA editing system

While the two-domain dCas13a-hADAR2d protein was programmed to edit reporter RNA, an important question is raised about flexibility of the fusion protein and how large its working parameter window is. As crRNA and pRNA fusion molecule was found to have a higher efficiency in directing RNA editing by dCas13a-hADAR2d, we proceeded to define its working parameter window using varying crRNA-pRNA fusion constructs.

To map out geometric constrains of the dCas13a-hADAR2d system, we designed and generated a series of crRNA-pRNA fusion constructs by placing the editing site residue (A) at variable positions along crRNA-pRNA fusion sequence. In practice, this was achieved with the editing site residue (A) being fixed, and the crRNA-pRNA sequence shifting from 3’ to 5’ (left to right) (Fig. 3*A*). Note this is possible due to the flexibility of the 3’ protospacer flanking site (PFS) for LshCas13a that permits either A, U, or C residue(7). As a result, we obtained eight different constructs, having the editing site residues (A) placed either inside pRNA region at sites +19, +15, +12, +6, or +2 bp, or inside crRNA region at sites −4, −5, or −9 bp, respectively, where numbers indicate the base position from crRNA/pRNA boundary (Fig. 3*A*). The ability of these constructs to direct site-specific RNA editing by dCas13a-hADAR2d was examined by the emission of green and red fluorescence in fission yeast cells carrying each construct. While the editing site residue (A) positioned inside crRNA region at −9, −5, and −4 displayed very little but detectable amount of green fluorescence, the signal increased as it moved into pRNA region, peaking around position +6 bp from the boundary (Fig. 3*B*). Our result revealed that optimized RNA editing activity by the dCas13a-hADAR2d system in *S. pombe* required the editing site residue to position ∼6 bp away from the crRNA region that interacts with dCas13a and target RNA.

**Figure 3.**
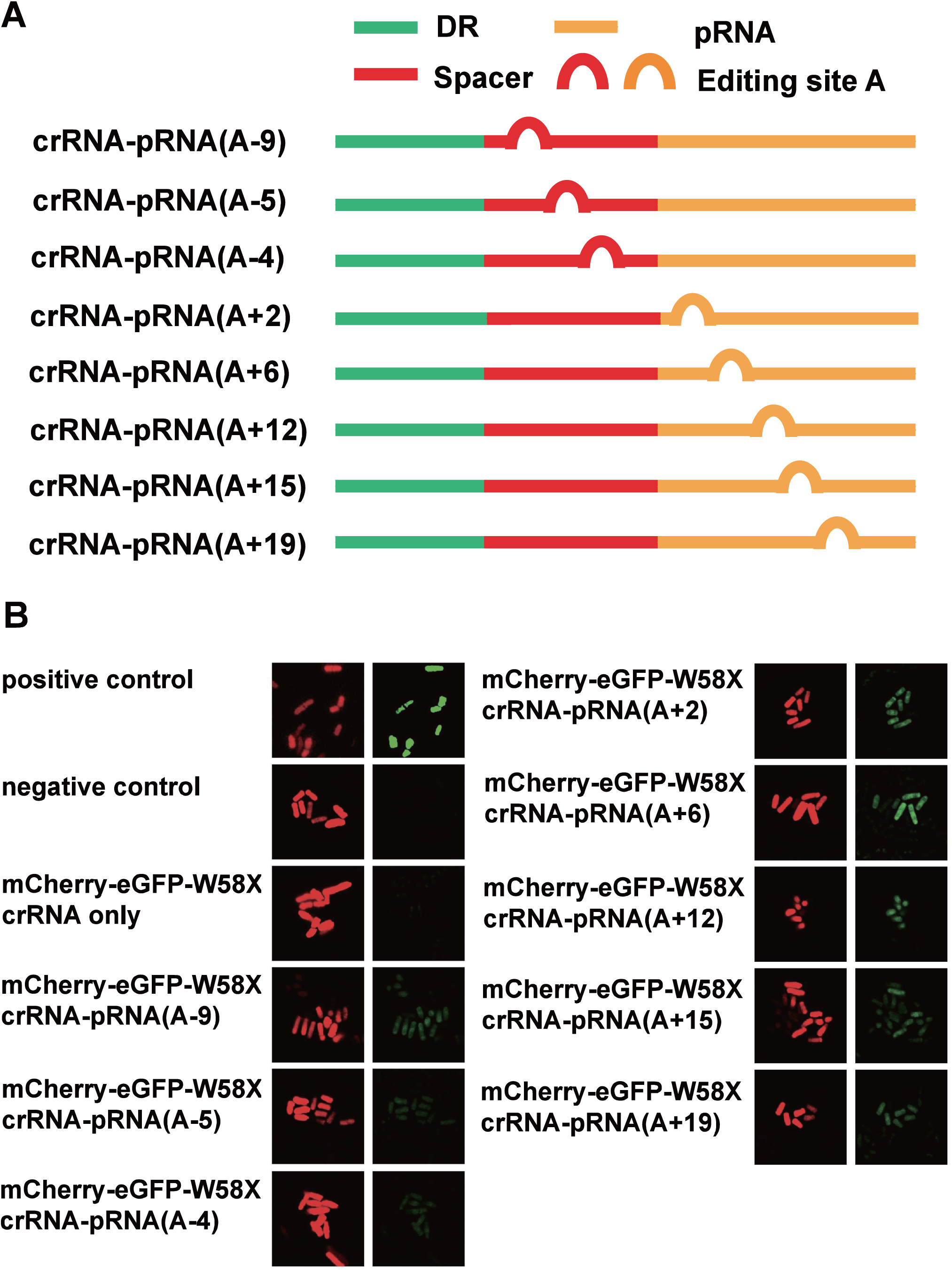
Defining working parameter window for dCas13a mediated RNA editing using fluorescent reporter. (A) Constructs of crRNA-pRNA fusion molecules as describe in Fig. 2*B*. The variable editing sites are achieved by shifting the crRNA-pRNA region along RNA sequence. The position of editing site residue (A) is marked (in bracket). ‘+’ and ‘-’ indicate editing sites inside pRNA region and crRNA region, respectively, and the numbers indicate the base position from crRNA/pRNA boundary. (B) Visualization of dCas13a-medicated RNA editing activity with editing site at variable position of crRNA-pRNA fusion molecules. Images were taken as described in Fig. 2*D*. Positive control, mCherry-eGFP in pDUAL-HFF1; negative control, mCherry-eGFP-W58X in pDUAL-HFF1.

### Length requirement of trans-acting pRNA for efficient RNA editing

We next determine the length requirement of pRNA for efficient mRNA editing by dCas13a-hADAR2d in *S. pombe*. Five crRNA-pRNA fusion constructs with varying pRNA length at 9, 11, 21, 31 and 37 bp were generated while the editing reside (A) was fixed within pRNA region at 6 bp from crRNA/pRNA boundary (Fig. 4*A*). We observed an increased green fluorescence signal as the pRNA length was extended from 9 to 37 bp, indicating better RNA editing activity was achieved with longer pRNA (Fig. 4*B*). The longer pRNA tended to form more stable dsRNA with target RNA, thus reflecting the requirement of dsRNA structure by the hADAR2d domain of the dCas13a-hADAR2d system in *S. pombe*.

**Figure 4.**
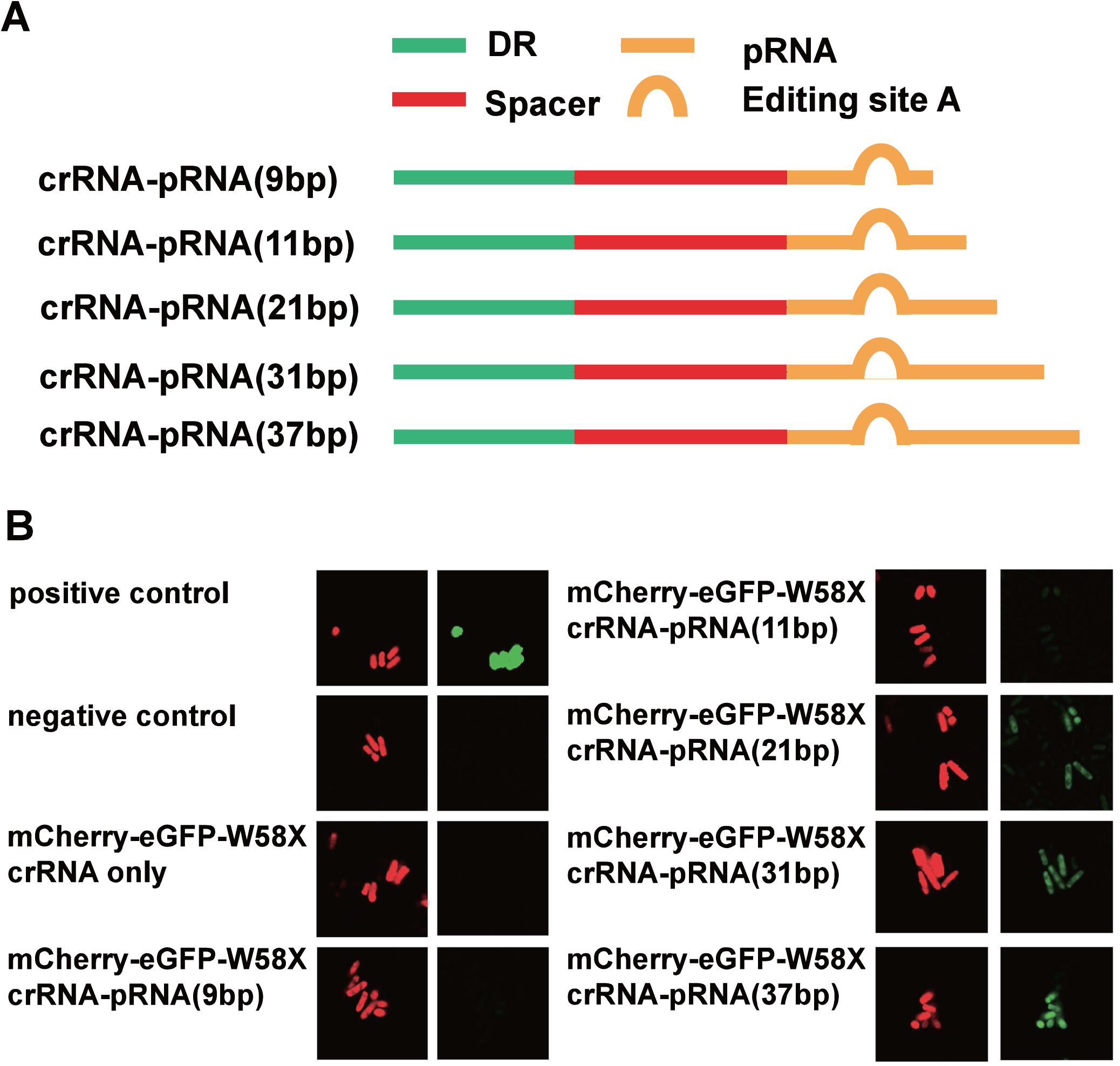
Defining length requirement of pRNA for efficient RNA editing by dCas13a-mediated system. (A) Constructs of crRNA-pRNA fusion molecules with variable length of pRNA. While the editing reside (A) is fixed in pRNA at 6 bp from crRNA/pRNA boundary, the length of pRNA varied at 9, 11, 21, 31, and 37 bp, as labeled in bracket. (B) Visualization of dCas13a-medicated RNA editing activity with variable length of pRNA. Images were taken as described in Fig. 2*D*. Positive control, mCherry-eGFP in pDUAL-HFF1; Negative control, mCherry-eGFP-W58X in pDUAL-HFF1.

### Application of dCas13a mediated RNA editing system on endogenous gene transcripts

To demonstrate the utility of our dCas13a-hADAR2d system in editing genes’ transcripts other than the fluorescence reporter, we designed and generated crRNA-pRNA constructs targeting a number of endogenous genes from *S. pombe* genome, i.e. *tdh1, act1, mel1, ade6, erp5, mug45*, and *nmt1*, which were randomly selected from *S. pombe* genome. These genes had an expression range from lowly expressed, e.g. *mel1* that has 0.069 mRNA molecules per cell in single-cell organism vegetative growth phase, to highly expressed, e.g. *tdh1* that has 560 mRNA molecules per cell during single-cell vegetative growth(36,37).

The crRNA-pRNA constructs that targeted editable UAG triplets within the genes’ transcripts, respectively (Table 1), were placed in the same plasmid construct (under *rrk1* promoter) as described (Fig. 2*B*). The plasmids were transformed into *S. pombe* strain with chromosomal dCas13a-hADAR2d gene. Site-specific RNA editing activity was detected by sequencing of reverse transcribed cDNA (*Materials and Methods*). For the nine sites targeted by our constructs, seven of them were found to have detectable RNA editing activity, with editing efficiency ranging from 4% to 59% (Table 1 and Fig. S2). Interestingly, although no editing activity was detected for *ade6* transcript at position 622A by Sanger sequencing (below detectable level), it was detected at two other sites, 1002A and 1160A, with 7% and 10% efficiency, respectively. The editing efficiency neither had apparent relationship with abundance of transcripts in cells, nor did with the G+C contents in the sequences flanking editing sites. Although *tdh1* and *nmt1* genes had comparable transcript abundance: 560 vs 340 mRNA molecules per cell(36), *tdh1* and *nmt1* had dramatically different editing efficiency at 59% and 4%, respectively. Overall, these results indicated that the dCas13a-hADAR2d system was applicable to a broad range of genes for RNA editing. At the same time, the observed varying activity amount target transcripts suggested the need for further refinement of system parameters in future applications.

**Table 1.**
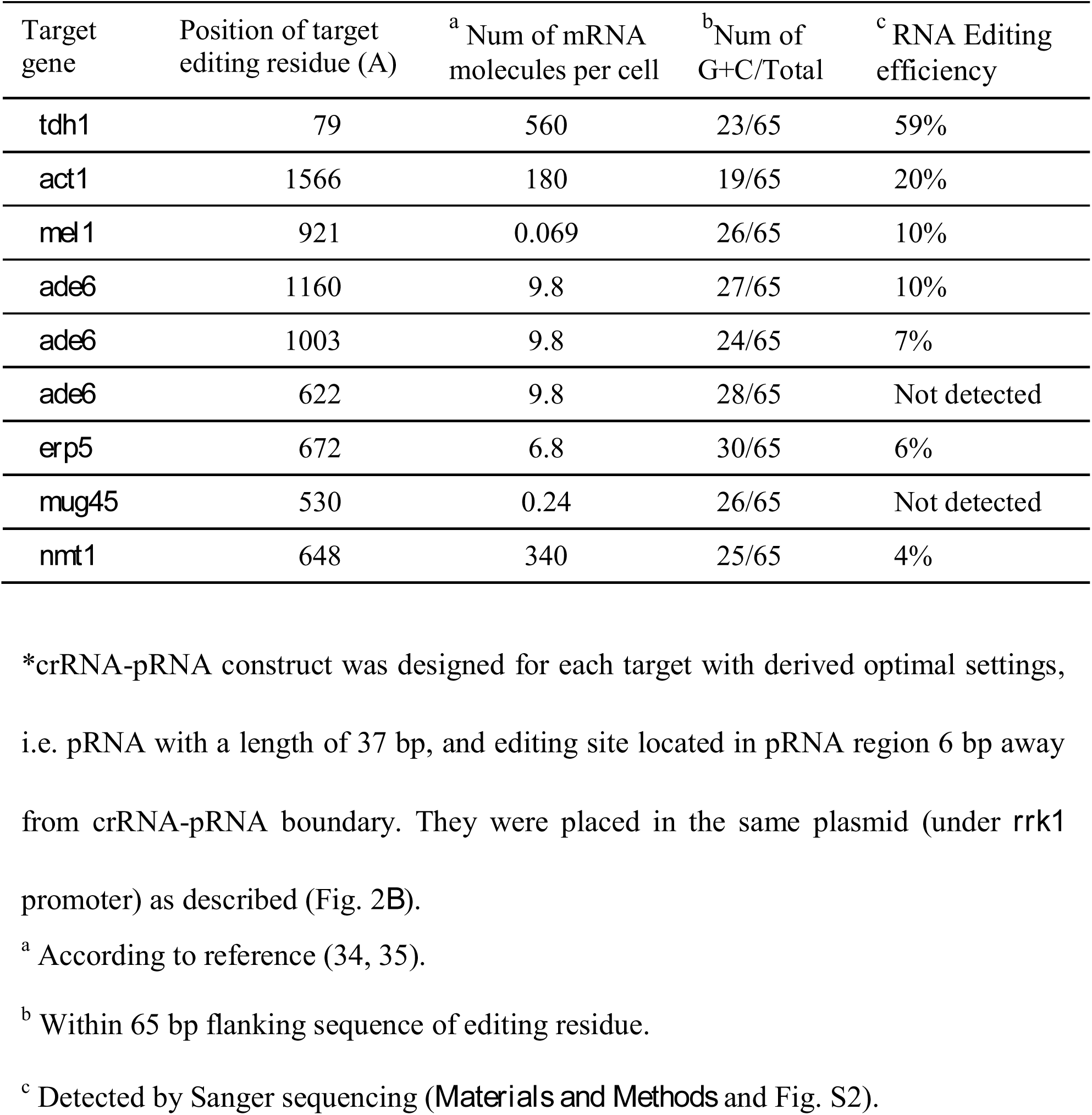
Editing of endogenous gene transcripts by dCas13a-hADAR2d in *S. pombe\**.

### Application of dCas13a mediated RNA editing system on restoring transposition of mutant TF1

To demonstrate the broad applicability of dCas13a mediated RNA editing system in *S. pombe*, we designed guides against four mutations in TF1 transpons: 836G→A(W99X), 1166 G→ A(W209X),3482 G→A(W981X) and 3851 G→A(W1104X). We transformed these mutant TF1 into *S. pombe* with dCas13a-hADAR2D and tested whether dCas13a mediated RNA editing system could correct the mutations and restore the TF1 transposition activity. Using guide RNAs containing 65nt, we were able to achieve 47.4% correction of 1166 G→A(W209X) and 68.9% correction of 3851 G→A(W1104X). No correction was detected for 836G→A(W99X) and 3482 G→A(W981X)(Fig 5*A*). We then tested the transponsition frequency of the mutant TF1 in *S. pombe*. For 836G→A(W99X), 1166 G→A(W209X) mutant TF1, no transposition was detected, and for 3482 G→A(W981X), 3851 G→A(W1104X) mutant TF1, 0.10% and 0.06% normalized transposition frequency were detected (Fig 5*B* and Table 2). When dCas13a mediated RNA editing system was introduced, the transposition frequency for 1166 G→A(W209X),3482 G→A(W981X) increased into 30% and 50% (Fig 5B and Table 2). While, no transposition was detected for 836G→A(W99X) TF1 with dCas13a mediated RNA editing system. The transposition frequency for 3482 G→A(W981X) TF1 was the same with or without dCas13a mediated RNA editing system.

## Discussion

CRISPR-Cas13a system has a single protein effector that contains HEPN domains and possesses two distinct RNAase activities for pre-crRNA processing and target RNA cleavage(7,8,10,38). We used the previously cloned Cas13a(7) from *Leptotrichia shahii* (LshCas13a) and successfully implemented it in the model organism *S. pombe*. LshCas13a was showed to target and knockdown endogenous gene transcripts in fission yeast cells. However, differently from what we expected basing on previous study *in vitro* and in *E. coli*(7,8,11), no collateral degradation of other non-specific RNA by activated Cas13a-crRNA complex was detected in fission yeast. The different behavior of LshCas13a in eukaryotic cells certainly warrants further investigation.

To implement LshCas13a in *S. pombe*, we applied *rrk1* promoter (transcribed by RNA Pol II) with a ribozyme cassette for *cis*-processing pre-crRNA. Similar system was previously used to implement CRISPR-Cas9 system in *S. pombe*(27). To meet the needs for both targeting RNA and editing applications of Cas13a, we made modifications to the existing cassette with the insertion of HDVR, and HHR1/HHR2 sequences (Fig. 2*B*). Such design was proved effective and efficient in generation of crRNA, or crRNA and pRNA molecules either separate or joined. Note the reported intrinsic RNase activity to process pre-crRNA by Cas13a could be utilized to simplify our system (8,38). However, in our design phase, we were concerned that the inactive LshCas13a (dCas13a) (to be used in constructing RNA editing complex) might have impaired pre-crRNA processing capability. One area of possible improvement for our system in future work is to test and utilize pre-crRNA processing function of dCas13a, for which could greatly reduce the complexity of the RNA editing toolset if successful.

On the basis of successful implementation of LshCas13a in *S. pombe*, we proceeded to design and engineer a toolset for targeted RNA editing, by tethering the inactive Cas13a (dCas13a) to the deaminase domain of human ADAR2 (hADAR2d). The dCas13a-hADAR2d fusion protein was reprogrammed to edit florescence reporter mRNA and endogenous genes’ transcripts. Although no study of targeted RNA editing tool was reported for yeast before, in mammalians cells a specially designed guide-RNA was introduced to recruit endogenous ADAR2 to edit specific mRNAs(39). Such approach was limited to cells expressing the endogenous RNA editing enzyme, i.e. ADAR. It also has a drawback of background activity from endogenous ADAR and relatively low binding affinity between guide-RNA and targets. Other works involved linking deaminase domain of human ADAR to a guide-RNA via either a SNAP-tag strategy or λN-boxB interaction(35,40-42). This category of methods depended on Watson and Crick base-paring for target recognition and binding, in addition to some needs for delivery of chemically modified guide-RNA (for SNAP-tag) to living cells(42-44). Our dCas13a-hADAR2d system provided an attractive alternative for RNA editing applications. Cas13a-crRNA and target-RNA binding is highly specific and tight. dCas13a (R1278A mutated LshCas3a) had an even stronger binding affinity towards its target RNA (K_D_ ∼ 7 nM) than the wild type LshCas13a (K_D_ ∼ 46 nM)(7). The high binding affinity made Cas13a suitable for a range of applications, for example, tagging for molecule localization and subcellular trafficking, RNA modifications, enrichment of specific RNA transcripts with their partners, etc(6,10,12).

While our data showed flexibility of dCas13a-hADAR2d in editing residue position, we were surprised to see some detectable, albeit very low activity of RNA editing for sites inside spacer region of crRNA (Fig. 3). The reported crystal structure of LshCas13a in complex with crRNA suggested that nucleotide bases 24-28 of the crRNA were exposed outside the LshCas13a structure(38). Further from the crystal structure of another close related Cas13a from *Lachnospiraceae bacterium* (LbaCas13a), the first 13 nucleotides (C1–G13) of the spacer in LbaCas13a/crRNA complex were well ordered and sequestered within LbaCas13a protein, whereas the rest 11 (G14–C24) were disordered and exposed in solvent(10). The disordered regions may suggest some possible opening for access by hADAR2d, resulting in residue RNA editing activity in spacer region.

The utilization of a different member of Cas13 family, Cas13b from *Prevotella sp. P5-125* (PspCas13b), to mediate RNA editing in human cell lines was reported(45). With an improved hyperactive hADAR2d variant (E488Q), the editing efficiency with PspCas13b/hADAR2d(E488Q) system was notably comparable to or better than the guide-RNA approaches discussed above(39,42-44). The performance of our dCas13a-hADAR2d system in *S. pombe* varied for different targets, with a RNA editing efficiency up to 59% for *tdh1* (Table 1), which is lower than the maximum performance of PspCas13b system in human cell lines. Note the vast difference existed in environment, design and configurations between the two systems. Notably, Cas13a-crRNA and Cas13b-crRNA complexes bind to their target RNA in opposite orientation(13,45). Thus our dCas13a-mediated RNA editing system has a completely different configuration from dCas13b-mediated system(45) (Fig. S4). In addition, factors, like gene transcript abundance, RNA secondary structure, and the molar ratio of dCas13a-hADAR2d to crRNA-pRNA or to target transcripts, contributed to the variation in editing efficiency. For example, an excessive amount of crRNA-pRNAs over dCas13a-hADAR2d molecules would prevent dCas13a-hADAR2d/crRNA-pRNA complex from acting on target RNAs by pre-occupying their proto-spacer sites on target RNA. It was also found the formation of Cas13a and crRNA complex was a pre-requisite for their recognition and binding of target RNA(10,38). However, Cas13a and crRNA complex may deny access by target RNA that has spatial obstacle due to their secondary structure and/or binding proteins. Further investigation is warranted to illustrate the system parameters underlying the specificity and efficiency of Cas13-medicated RNA interference system.

While we have shown that Cas13a can be implemented and repurposed for RNA-interference and RNA editing tasks in *S. pombe*, we foresee that it can be improved and expanded to many other functions, addressing the fundamental questions regarding to modulation of gene functions at RNA levels using the model system of fission yeast. To make the CRISPR-Cas13 related tool more effective, it can be enhanced with new functionality. For example, our dCas13a-hADAR2d system in *S. pombe* can be further improved by simply using the E488Q variant of hADAR2d. Or we can look to expand RNA editing activity by replacing hADAR2d with RNA editing enzyme of different specificity, such as the APOBEC family of enzymes that catalyze cytidine to uridine conversion. While molecular tools manipulating RNA sequences are limited, this RNA manipulation platform is specially valuable for perturbing model organisms like *S. pombe*. Our implementation of CRISPR-13a system in fission yeast served as a stepping-stone for much promised, new family of toolsets that target gene transcripts for a wide range of research and biotechnological applications.

## Acknowledgements

This work was supported in part by grants from the Ministry of Agriculture of China [2016ZX08010-002]; the National Natural Science Foundation of China [nos. 31401128, 31571310, 31771412]; and by Special Fund for Strategic Pilot Technology Chinese Academy of Sciences [XDA08020104].

## Author contributions

X.L. conceived the project, designed experiments, and wrote the manuscript. S.Y. and P.H. directed/advised on the study. X.J. performed experiments, analyzed data, and prepared the manuscript. B.X., L.C., H.Q., Y.J. and N.B.Z. performed experiments and helped with data analysis.

