## Supplementary Materials for "Implementation of CRISPR-Cas13a system in fission yeast and its repurposing for precise RNA editing"

**Figure S1**

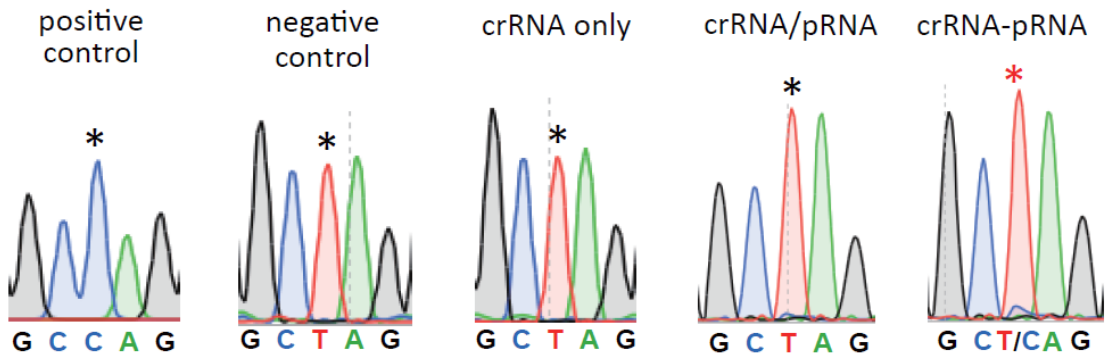

**Figure S1.** Electropherograms from RT-PCR products of *S. pombe* with chromosomal dCas13a-ADAR2d gene carrying different plasmids construct (*SI Methods* and Table S5). Positive control, mCherry-eGFP in pDUAL-HFF1; negative control, mCherry-eGFP-W58X in pDUAL-HFF1.

**Figure S2**

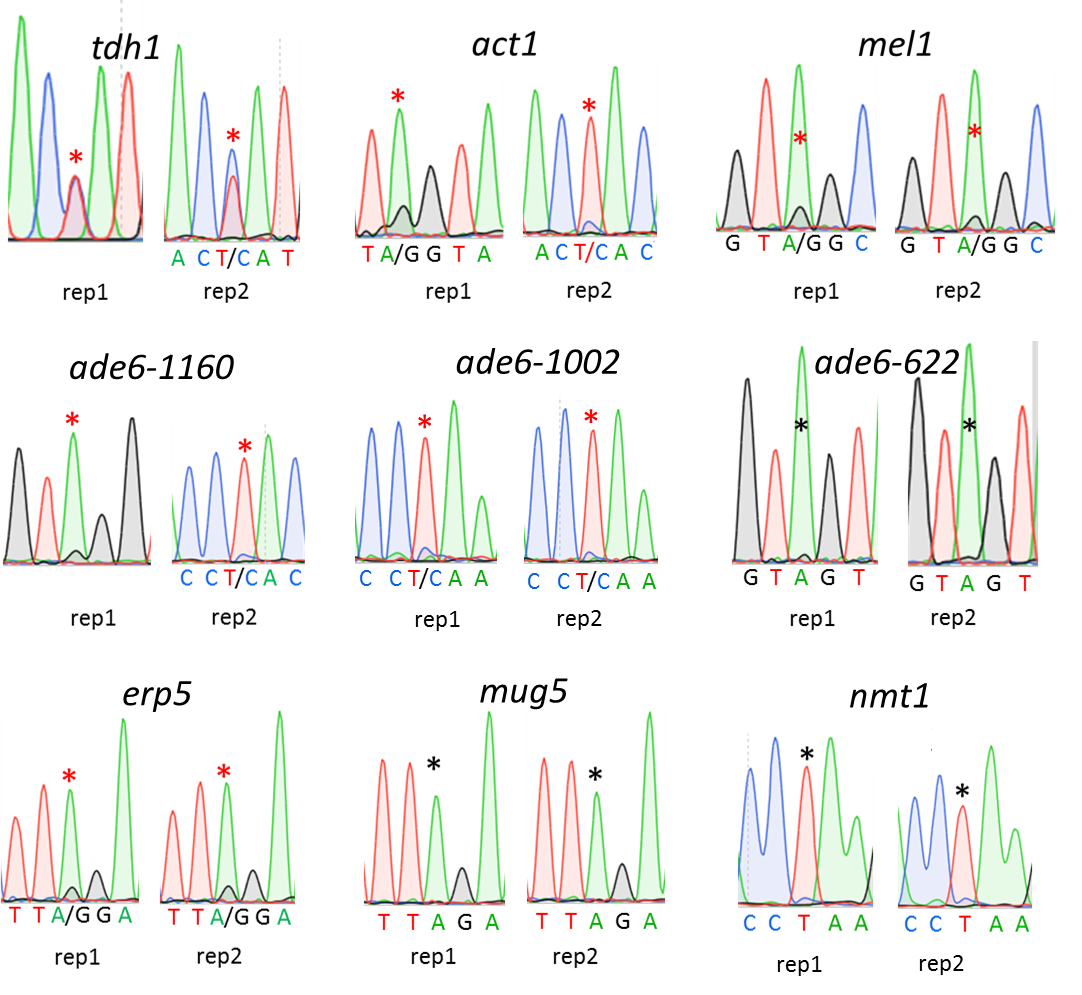

**Figure S2.** Electropherograms from RT-PCR products of *S. pombe* carrying plasmids with episomal dCas13a-ADAR2d gene and different crRNA-pRNA construct targeting endogenous gene transcripts (*SI Methods* and Table S5). crRNA-pRNA construct was designed for each target with derived optimal settings, i.e. pRNA with a length of 37 bp, and editing site located in pRNA region 6 bp away from crRNA-pRNA boundary. They were placed in the same plasmid (under *rrk1* promoter) as described (Fig. 2*B*)

**Figure S3**

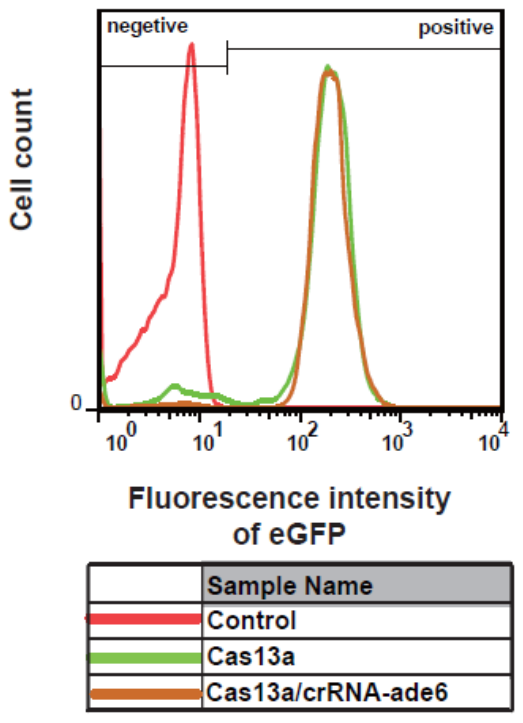

**Figure S3.** Distribution of florescence intensity in *S. pombe* cells. Control, *S. pombe* (FY7652); Cas13a, *S. pombe* strain (with chromosomal eGFP gene) transformed with pDUAL-HFF1-Cas13a; Cas13a/crRNA-ade6, *S. pombe* strain (with chromosomal eGFP gene) transformed with pDUAL-HFF1-Cas13a-rrk1-crRNA-ade6.

**Figure S4**

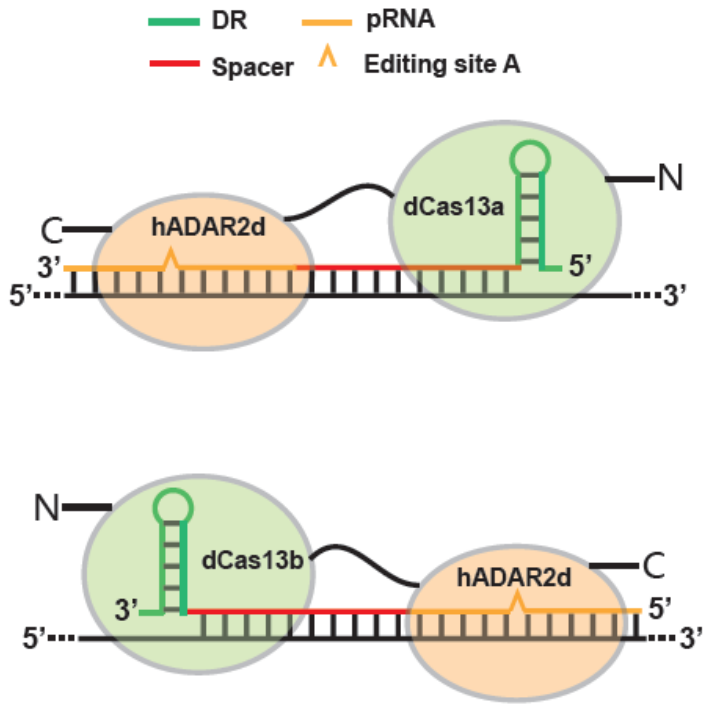

**Figure S4.** Schematic comparison of RNA editing constructs between our dCas13a-hADAR2d and that of dCas13b. Note dCas13a-mediated RNA editing system has a completely different configuration from dCas13b-mediated system.

**Figure S5**

atggggaacctgttcggacacaagcggtggtacgaggtgcgagataagaaagacttcaagatcaagcgaaaggtgaaagtcaagcggaattatgacgggaacaagtacattctgaacatcaacgaaaacaacaacaaagagaagatcgacaacaacaagttcatcagaaagtacatcaactacaagaaaaacgataatattctgaaggagttcactaggaaatttcatgcaggaaatatcctgttcaaactgaagggcaaagaagggatcattagaattgagaataacgacgacttcctggagacagaggaagtggtgctgtatatcgaggcctacggcaagagcgagaagctgaaagcactggggatcacaaagaaaaagatcattgacgaggccatcaggcagggcattactaaggacgataaaaagatcgagatcaagcgacaggagaacgaggaagagatcgaaattgacatccgggatgagtacactaataagaccctgaacgactgctccatcattctgcgcatcattgaaaacgatgaactggagacaaagaagagcatctacgagatcttcaagaacatcaatatgagcctgtataagatcatcgagaagatcatcgaaaacgagacagaaaaggtgtttgaaaatagatactacgaagagcacctgagggagaagctgctgaaagacgataagattgacgtgatcctgaccaacttcatggaaatccgggagaagatcaagtctaatctggagatcctgggcttcgtgaagttttacctgaacgtcggcggggacaaaaagaaaagtaaaaataagaaaatgctggtggaaaagattctgaacatcaatgtggatctgaccgtcgaggacattgccgatttcgtgatcaaggagctggaattttggaacatcacaaagcgcattgagaaagtgaagaaagtcaataacgagttcctggagaagcggagaaatcggacatacatcaagtcctatgtgctgctggacaagcacgaaaagtttaaaatcgagagagaaaacaagaaggataagatcgtgaagttctttgtcgagaacattaagaacaactctatcaaggaaaagattgagaagatcctggctgagttcaagatcgacgagctgattaagaaactggagaaggaactgaagaaagggaactgtgataccgagatcttcggaatctttaagaagcattacaaggtgaacttcgacagcaagaaattttccaagaaatctgatgaagagaaggagctgtataagatcatctacagatacctgaagggcagaattgaaaaaatcctggtgaacgagcagaaggtcagactgaagaaaatggagaagatcgagatcgaaaagattctgaatgaaagtatcctgtcagagaaaattctgaagagagtgaaacagtatacactggagcacattatgtacctggggaagctgaggcataacgacatcgatatgaccacagtgaatactgacgatttcagccgcctgcacgccaaggaagagctggacctggaactgatcaccttctttgccagcacaaatatggagctgaacaagatcttttcccgagaaaacatcaacaacgacgagaacatcgatttctttggaggcgaccgggagaagaactatgtgctggataagaaaatcctgaatagtaagatcaagatcatccgcgacctggatttcatcgataacaagaacaacatcacaaacaacttcattcgaaagtttacaaagatcggcactaatgaaaggaaccgcatcctgcatgccatttccaaagagagggacctgcaggggactcaggacgattacaacaaagtgatcaacatcattcagaatctgaagatctccgatgaagaggtgagcaaagctctgaacctggacgtggtctttaaggacaagaaaaacatcatcacaaagatcaatgacatcaagatctctgaagagaacaacaacgatatcaagtatctgcccagcttcagcaaagtgctgcccgaaatcctgaacctgtaccgcaacaatcccaagaatgagccttttgacacaatcgagactgaaaaaattgtgctgaacgctctgatctacgtcaataaggagctgtataagaaactgatcctggaggacgatctggaagagaacgagtccaagaatatcttcctgcaggaactgaagaaaaccctgggcaacattgacgaaatcgatgagaacatcatcgagaactactacaagaacgcacagatttctgccagtaaggggaacaacaaggcaatcaagaaatatcagaagaaagtgatcgagtgctacattggatatctgcgcaaaaactacgaagagctgttcgacttttcagacttcaagatgaacatccaggaaatcaagaaacagattaaggacatcaacgataacaagacttatgagcggatcaccgtgaaaaccagcgacaaaaccattgtcatcaacgacgatttcgagtacatcatttctatctttgcactgctgaacagtaatgccgtgattaataagatccgaaacagattcttcgccaccagcgtgtggctgaacacctcagaataccagaatatcattgacatcctggatgagattatgcagctgaataccctgcggaacgaatgcatcacagagaactggaatctgaacctggaagagttcattcagaagatgaaagagatcgaaaaggatttcgacgacttcaagatccagactaagaaagaaatcttcaacaactactacgaggacatcaagaacaacattctgaccgagtttaaagacgatatcaacggctgtgatgtgctggaaaagaaactggagaagattgtcatcttcgacgatgaaaccaagttcgagatcgacaagaaatccaacatcctgcaggatgaacagagaaagctgtctaacatcaacaagaaggacctgaagaaaaaggtggatcagtatatcaaggacaaagatcaggagatcaagtctaaaatcctgtgcaggatcattttcaacagtgactttctgaaaaagtacaagaaggaaatcgacaatctgattgaggatatggagtctgaaaatgagaacaagttccaggagatctactatcccaaggaacggaagaacgagctgtatatctacaaaaagaatctgttcctgaacatcggaaatcctaactttgacaagatctacggcctgattagcaacgacatcaagatggccgatgctaaattcctgtttaatatcgatggaaagaacatcagaaagaacaaaatcagtgagatcgacgctattctgaagaatctgaacgataaactgaacggctactcaaaggaatacaaggagaagtacatcaaaaagctgaaggagaacgacgatttctttgcaaagaacatccagaataagaactacaaatccttcgaaaaggactataaccgcgtgtctgagtacaaaaagattcgagatctggtcgagttcaactatctgaacaaaatcgagtcctacctgattgacatcaactggaagctggctattcagatggcaagattcgaaagggatatgcactatatcgtgaatggactgagggagctgggcatcattaagctgtcaggctataacaccgggatcagcagggcatacccaaagcgcaatggaagcgacggcttttacactaccacagcctactacaagttctttgatgaagagtcctacaagaagttcgagaagatttgctacgggtttggaatcgacctgagcgaaaattccgagatcaacaagcctgaaaatgagagcattgctaactatatctcccatttctacatcgtgagaaatccatttgccgactacagtattgctgagcagatcgatcgggtgagcaacctgctgtcatatagcacacgctacaacaattcaacttatgccagcgtgttcgaagtctttaaaaaggacgtgaatctggactacgatgagctgaaaaagaaattcaaactgatcggcaacaatgatattctggagcgcctgatgaagcccaagaaagtgagcgtcctggaactggagtcctacaacagtgactacattaagaatctgatcattgaactgctgaccaaaatcgagaatactaacgataccctgaaatctggttctgaaactcctggtacttctgaatctgctactcctgaattgcacttggatcagacgccatctcgccagcctattcccagtgagggtcttcagctgcatttaccgcaggttttagctgacgctgtctcacgcctggtcctgggtaagtttggtgacctgaccgacaacttctcctcccctcacgctcgcagaaaagtgctggctggagtcgtcatgacaacaggcacagatgttaaagatgccaaggtgataagtgtttctacaggaacaaaatgtattaatggtgaatacatgagtgatcgtggccttgcattaaatgactgccatgcagaaataatatctcggagatccttgctcagatttctttatacacaacttgagctttacttaaataacaaagatgatcaaaaaagatccatctttcagaaatcagagcgaggggggtttaggctgaaggagaatgtccagtttcatctgtacatcagcacctctccctgtggagatgccagaatcttctcaccacatgagccaatcctggaagaaccagcagatagacacccaaatcgtaaagcaagaggacagctacggaccaaaatagagtctggtgaggggacgattccagtgcgctccaatgcgagcatccaaacgtgggacggggtgctgcaaggggagcggctgctcaccatgtcctgcagtgacaagattgcacgctggaacgtggtgggcatccagggatccctgctcagcattttcgtggagcccatttacttctcgagcatcatcctgggcagcctttaccacggggaccacctttccagggccatgtaccagcggatctccaacatagaggacctgccacctctctacaccctcaacaagcctttgctcagtggcatcagcaatgcagaagcacggcagccagggaaggcccccaacttcagtgtcaactggacggtaggcgactccgctattgaggtcatcaacgccacgactgggaaggatgagctgggccgcgcgtcccgcctgtgtaagcacgcgttgtactgtcgctggatgcgtgtgcacggcaaggttccctcccacttactacgctccaagattaccaagcccaacgtgtaccatgagtccaagctggcggcaaaggagtaccaggccgccaaggcgcgtctgttcacagccttcatcaaggcggggctgggggcctgggtggagaagcccaccgagcaggaccagttctcactcacgccctga

**Figure S5.** Gene sequences of dCas13a-hADAR2d in the context of pDUAL-HFF1 plasmid, under the promoter of *nmt1* and ADH1 terminator. Nucleotides for R1278A mutant of Cas13a protein, green background; XTEN linker sequence with codon optimized for expression in *S. pombe*: magenta background.

**Figure S6**

ATGGTGAGCAAGGGCGAGGAGGATAACATGGCCATCATCAAGGAGTTCATGCGCTTCAAGGTGCACATGGAGGGCTCCGTGAACGGCCACGAGTTCGAGATCGAGGGCGAGGGCGAGGGCCGCCCCTACGAGGGCACCCAGACCGCCAAGCTGAAGGTGACCAAGGGTGGCCCCCTGCCCTTCGCCTGGGACATCCTGTCCCCTCAGTTCATGTACGGCTCCAAGGCCTACGTGAAGCACCCCGCCGACATCCCCGACTACTTGAAGCTGTCCTTCCCCGAGGGCTTCAAGTGGGAGCGCGTGATGAACTTCGAGGACGGCGGCGTGGTGACCGTGACCCAGGACTCCTCCCTGCAGGACGGCGAGTTCATCTACAAGGTGAAGCTGCGCGGCACCAACTTCCCCTCCGACGGCCCCGTAATGCAGAAGAAGACCATGGGCTGGGAGGCCTCCTCCGAGCGGATGTACCCCGAGGACGGCGCCCTGAAGGGCGAGATCAAGCAGAGGCTGAAGCTGAAGGACGGCGGCCACTACGACGCTGAGGTCAAGACCACCTACAAGGCCAAGAAGCCCGTGCAGCTGCCCGGCGCCTACAACGTCAACATCAAGTTGGACATCACCTCCCACAACGAGGACTACACCATCGTGGAACAGTACGAACGCGCCGAGGGCCGCCACTCCACCGGCGGCATGGACGAGCTGTACAAGGAATTCtggaacatggcctcccgaggagcccagacagctgcagccacagctccccgtatcaagaaatttgccatctatcgatgggacccagacaaggctggagacaaacctcatTCTAGAGTGAGCAAGGGCGAGGAGCTGTTCACCGGGGTGGTGCCCATCCTGGTCGAGCTGGACGGCGACGTAAACGGCCACAAGTTCAGCGTGTCCGGCGAGGGCGAGGGCGATGCCACCTACGGCAAGCTGACCCTGAAGTTCATCTGCACCACCGGCAAGCTGCCCGTGCCCTGGCCCACCCTCGTGACCACCCTGACCTACGGCGTGCAGTGCTTCAGCCGCTACCCCGACCACATGAAGCAGCACGACTTCTTCAAGTCCGCCATGCCCGAAGGCTACGTCCAGGAGCGCACCATCTTCTTCAAGGACGACGGCAACTACAAGACCCGCGCCGAGGTGAAGTTCGAGGGCGACACCCTGGTGAACCGCATCGAGCTGAAGGGCATCGACTTCAAGGAGGACGGCAACATCCTGGGGCACAAGCTGGAGTACAACTACAACAGCCACAACGTCTATATCATGGCCGACAAGCAGAAGAACGGCATCAAGGTGAACTTCAAGATCCGCCACAACATCGAGGACGGCAGCGTGCAGCTCGCCGACCACTACCAGCAGAACACCCCCATCGGCGACGGCCCCGTGCTGCTGCCCGACAACCACTACCTGAGCACCCAGTCCGCCCTGAGCAAAGACCCCAACGAGAAGCGCGATCACATGGTCCTGCTGGAGTTCGTGACCGCCGCCGGGATCACTCTCGGCATGGACGAGCTGTACAAGTCCGGATAA

**Figure S6.** Gene sequences of mCherry-eGFP-W58X in the context of pDUAL-HFF1 plasmid, under the promoter of *nmt1* and ADH1 terminator. Nucleotide for the W58X mutant of eGFP protein, red background.

**Figure S7**

TTTTGCTTATGTTGGTGGTAGTTGGCATGCGTAGACTGATGACTAGTCAGCAAGGAGCGTAGAACAGTCACACTCGTTATATATGTGCTTCCAAGAAAACTCAAGAATTTACCATTAGCAAACACTTTTTTGAAATGTTAGACATTTAAATGACGAAGGCATATAGAAGCTTTGAATAGGTGTTGTAAAGTGTTGATTTATGTGACGCTGAGGGTGCGCATGAAAGGAATGTTGGGTCACGATTATTAAACAGTTTGCTAGCTTGGACACTTGAGTATTGGAAGTTGTTGAATTCTAAAAAACTTTCAGTTGATTTGAATAGTTGCTGTTGCCAAAAAACATAACCTGTACCGAAGAAccaccccaatatcgaaggggactaaaacAGAAGAGCTGAATTCAGCTCTTCA**GGCCGGCATGGTCCCAGCCTCCTCGCTGGCGCCGGCTGGGCAACATGCTTCGGCATGGCGAATGGGAC**agagacctgaattcaggtctcaCCTGTCACCGGATGTGCTTTCCGGTCTGATGAGTCCGTGAGGACGAAACAGG

**Figure S7.** crRNA-pRNA cassete sequence with *rrk1* promoter. *rrk1*promoter and leader RNA, capital letters; DR, lowercase letters; *Bsp*QI place holder for crRNA, underlined capital letters; HDVR, bold letters; *Bsa*I placer holder for pRNA, underlined lowercase letters; HHR, wave underlined captical letters.

**Table S1.** List of main strains and plasmids used in this study.

| Strains and plasmids | Characteristics | Source/Reference |
| --- | --- | --- |
| **Strains** | | |
| *E.coli* [DH5α](http://www.baidu.com/link?url=_2cqaHxXn_OKbZfgxUEUswAADXyKK09VwEG8ZiomAceKhUSCMxNulzbJr5ie7KjuzMUnC8yk6ZDUVYta7zUVX0HPMFVFV892zvJbyLgOgye2VtsFxWEMMjAyXoFV-l4e4HRZN4eVDAVnQ6A0IwU5-F2YvJzb4AI4NnlG3IT3ba49B0aVXWKRpjLVdZX-QcE97cgbbO3wKRwD22R2Vx5d_q&wd=&eqid=e461002d00003cd1000000035a096904) | *F- eNDA1 glnV44 thi-1 recA1 relA1 gyrA96 deoR nupG*  *Φ80dlacZΔM15 Δ(lacZYA-argF)U169, hsdR17 (rK-mK), λ–* | Takara Biotechnology Co.,Ltd. |
| *S.pombe* FY7652 | *h- leu1-32 ura4-D18* | National Bio Resource Project |
| **Plasmids** | | |
| pBluescript II KS(+) | *E.coli* , *ColE1 ori*, F1 *ori*, Ap ^r^ |  |
| pDUAL-HFF1 | *E.coli* -*S.pombe* shutting vector, *ars1*, *ori*, Ap ^r^ , *ura4* | RIKEN BRC (RDB:6179) |
| pC001 | *E.coli* , *ColE1 ori*, F1 *ori*, Ap ^r^, Cas13a, | Addgene: 79150 |
| pEGFP-N1-FLAG | *E.coli,* eGFP | Addgene: 60360 |
| pmCherry Paxillin | *E.coli,* mCherry | Addgene: 50526 |
| pDUAL-HFF1-eGFP | *FY7652,* eGFP/*nmt1* | This study |
| pDUAL-HFF1-mCherry-eGFP | *FY7652,* mCherry, eGFP/*nmt1* | This study |
| pDUAL-HFF1-mCherry-eGFP-W58X | *FY7652,* mCherry/*nmt1* | This study |
| pDUAL-HFF1-Cas13a | *FY7652,* Cas13a/*nmt1* | This study |
| pDUAL-HFF1-dCas13a-hADAR2d | *FY7652,* dCas13a-hADAR2d/*nmt1* | This study |
| pKS-rrk1-crRNA-backbone | *E.coli,* crRNA-pRNA expression cassete, Amp^r^ | This study |

**TABLE S2.** Intermediate plasmids for reporter and crRNA/pRNA constructs.

| **Intermediate plasmids** | **Primers** | **Cloning sites** | **Original plasmids** | **Source** |
| --- | --- | --- | --- | --- |
| pKS-mCherry-linker | mCherry-P5  mCherry-linker-P3  mCherry-linker-P5  linker-P3 | *Kpn*I/*Xba*I | pBluescript II KS(+) | This study |
| pKS-mCherry-eGFP | eGFP-P5  eGFP-P3 | *Xba*I/*Bgl*II | pKS-mCherry-linker | This study |
| pKS-mCherry-eGFP-W58X | eGFP-P5  eEGFP-mut-P3  eEGFP-mut-P5  eGFP-P3 | *Xba*I/*Bgl*II | pKS-mCherry-linker | This study |
| pKS-rrk1-crRNA-control | crRNA-eGFP-P5  crRNA-eGFP-P3 | *BspQ*I | pKS-rrk-crRNA-backbone | This study |
| pKS-rrk1-crRNA-pRNA-seperate | crRNA-eGFP-P5  crRNA-eGFP-P3 | crRNA/*BspQ*I  pRNA/*Bsa*I | pKS-rrk-crRNA-backbone | This study |
| pKS-rrk1-crRNA-pRNA-fusion | crRNA-L-P5  crRNA-L-P3 | *BspQ*I | pKS-rrk-crRNA-backbone | This study |
| pKS-rrk1-crRNA-pRNA-6-37 | crRNA-eGFP-fusion-6-37-p5  crRNA-eGFP-fusion-6-37-p3 | *BspQ*I | pKS-rrk-crRNA-backbone | This study |
| pKS-rrk1-crRNA-pRNA-6-39 | crRNA-eGFP-fusion-6-39-p5  crRNA-eGFP-fusion-6-39-p3 | *BspQ*I | pKS-rrk-crRNA-backbone | This study |
| pKS-rrk1-crRNA-pRNA-6-49 | crRNA-eGFP-fusion-6-49-p5  crRNA-eGFP-fusion-6-49-p3 | *BspQ*I | pKS-rrk-crRNA-backbone | This study |
| pKS-rrk1-crRNA-pRNA-6-59 | crRNA-eGFP-fusion-6-59-p5  crRNA-eGFP-fusion-6-59-p3 | *BspQ*I | pKS-rrk-crRNA-backbone | This study |
| pKS-rrk1-crRNA-pRNA-(-9)-65 | crRNA-eGFP-fusion-(-9)-65-p5  crRNA-eGFP-fusion-(-9)-65-p3 | *BspQ*I | pKS-rrk-crRNA-backbone | This study |
| pKS-rrk1-crRNA-pRNA-(-5)-65 | crRNA-eGFP-fusion-(-5)-65-p5  crRNA-eGFP-fusion-(-5)-65-p3 | *BspQ*I | pKS-rrk-crRNA-backbone | This study |
| pKS-rrk1-crRNA-pRNA-(-4)-65 | crRNA-eGFP-fusion-(-4)-65-p5  crRNA-eGFP-fusion-(-4)-65-p3 | *BspQ*I | pKS-rrk-crRNA-backbone | This study |
| pKS-rrk1-crRNA-pRNA-2-65 | crRNA-eGFP-fusion-(2)-65-p5  crRNA-eGFP-fusion-(2)-65-p3 | *BspQ*I | pKS-rrk-crRNA-backbone | This study |
| pKS-rrk1-crRNA-pRNA-12-65 | crRNA-eGFP-fusion-(12)-65-p5  crRNA-eGFP-fusion-(12)-65-p3 | *BspQ*I | pKS-rrk-crRNA-backbone | This study |
| pKS-rrk1-crRNA-pRNA-15-65 | crRNA-eGFP-fusion-(15)-65-p5  crRNA-eGFP-fusion-(15)-65-p3 | *BspQ*I | pKS-rrk-crRNA-backbone | This study |
| pKS-rrk1-crRNA-pRNA-19-65 | crRNA-eGFP-fusion-(19)-65-p5  crRNA-eGFP-fusion-(19)-65-p3 | *BspQ*I | pKS-rrk-crRNA-backbone | This study |
| pKS-rrk1-crRNA-pRNA-act1-1566 | crRNA-act1-1566-p5  crRNA-act1-1566-p3 | *BspQ*I | pKS-rrk-crRNA-backbone | This study |
| pKS-rrk1-crRNA-pRNA-ade6-622 | crRNA-ade6-622-p5  crRNA-ade6-622-p3 | *BspQ*I | pKS-rrk-crRNA-backbone | This study |
| pKS-rrk1-crRNA-pRNA-ade6-1002 | crRNA-ade6-1003-p5  crRNA-ade6-1003-p3 | *BspQ*I | pKS-rrk-crRNA-backbone | This study |
| pKS-rrk1-crRNA-pRNA-ade6-1160 | crRNA-ade6-1160-p5  crRNA-ade6-1160-p3 | *BspQ*I | pKS-rrk-crRNA-backbone | This study |
| pKS-rrk1-crRNA-pRNA-erp5-672 | crRNA-erp5-672-p5  crRNA-erp5-672-p3 | *BspQ*I | pKS-rrk-crRNA-backbone | This study |
| pKS-rrk1-crRNA-pRNA-mel1-921 | crRNA-mel1-921-p5  crRNA-mel1-921-p3 | *BspQ*I | pKS-rrk-crRNA-backbone | This study |
| pKS-rrk1-crRNA-pRNA-mug45 | crRNA-mug45-530-p5  crRNA-mug45-530-p3 | *BspQ*I | pKS-rrk-crRNA-backbone | This study |
| pKS-rrk1-crRNA-pRNA-nmt1-648 | crRNA-nmt1-648-p5  crRNA-nmt1-648-p3 | *BspQ*I | pKS-rrk-crRNA-backbone | This study |
| pKS-rrk1-crRNA-pRNA-tdh1-79 | crRNA-tdh1-79-p5  crRNA-tdh1-79-p3 | *BspQ*I | pKS-rrk-crRNA-backbone | This study |

**TABLE S3.** Two endogenous genes from *S. pombe* with its transcripts targeted by crRNA constructs.

| **Gene name** | **Gene ID** | **Transcript Size** | **protospacer**  **location** | **Target sequence by crRNA** |
| --- | --- | --- | --- | --- |
| *ade6* | SPCC1322.13 | 1745 | 1130-1194 | ATTTCTGATTCACCTCAAGAATGTGAACGTaGGTATCAGATGCTTCTTGACGTCAAAGATCCTGT |
| *tdh1* | SPBC32F12.11 | 1518 | 49-113 | GCATCGCTTCTGTATAGATCATTCATCCATAGTATTGATTTACACTTGATTCAAAATGGCAATTC |

**TABLE S4.** Constructs for knocking down endogenous gene.

| **Plasmid Name** | **Genes/Promoters** | **crRNA-pRNA primers** | **Original plasmid** | **Targeting gene** | **Source** |
| --- | --- | --- | --- | --- | --- |
| pDUAL-HFF-Cas13a  -rrk1-crRNA-ade6-1160 | *cas13a*/*nmt1*  crRNA/*rrk1* | crRNA-ade6-1160-p5  crRNA-ade6-1160-p3 | pDUAL-HFF1 | *ade6* | This study |
| pDUAL-HFF-Cas13a  -rrk1-crRNA-tdh1-79 | *cas13a*/*nmt1*  crRNA/*rrk1* | crRNA-tdh1-79-p5  crRNA-tdh1-79-p3 | pDUAL-HFF1 | *tdh1* | This study |

**TABLE S5.** crRNA-pRNA constructs for editing of reporter gene mCherry-eGFP-W58X.

| **Plasmid Name** | **Genes/Promoters** | **crRNA-pRNA primers** | **crRNA-pRNA**  **length (nt)** | **Editing site** | **Source** |
| --- | --- | --- | --- | --- | --- |
| pDUAL-HFF-mCherry-eGFP-W58X  -rrk1-crRNA-control | mCherry-eGFP-W58X /*nmt1*  crRNA/*rrk1* | crRNA-eGFP-P5  crRNA-eGFP-P3 | 28 |  | This study |
| pDUAL-HFF-mCherry-eGFP-W58X  -rrk1-crRNA-pRNA-seperate | mCherry-eGFP-W58X /*nmt1*  crRNA/pRNA/*rrk1* | crRNA-eGFP-P5  crRNA-eGFP-P3  pRNA-HH-P5  pRNA-HH-P3 | 28+37 | 6 | This study |
| pDUAL-HFF-mCherry-eGFP-W58X  -rrk1-crRNA-pRNA-fusion | mCherry-eGFP-W58X /*nmt1*  crRNA-pRNA/*rrk1* | crRNA-L-P5  crRNA-L-P3 | 65 | 6 | This study |
| pDUAL-HFF-mCherry-eGFP-W58X  -rrk1-crRNA-6-37 | mCherry-eGFP-W58X /*nmt1*  crRNA-pRNA(9bp)/*rrk1* | crRNA-eGFP-fusion-6-37-p5  crRNA-eGFP-fusion-6-37-p3 | 28+9 | 6 | This study |
| pDUAL-HFF-mCherry-eGFP-W58X  -rrk1-crRNA-6-39 | mCherry-eGFP-W58X /*nmt1*  crRNA-pRNA(11bp)/*rrk1* | crRNA-eGFP-fusion-6-39-p5  crRNA-eGFP-fusion-6-39-p3 | 28+11 | 6 | This study |
| pDUAL-HFF-mCherry-eGFP-W58X  -rrk1-crRNA-6-49 | mCherry-eGFP-W58X /*nmt1*  crRNA-pRNA(21bp)/*rrk1* | crRNA-eGFP-fusion-6-49-p5  crRNA-eGFP-fusion-6-49-p3 | 28+21 | 6 | This study |
| pDUAL-HFF-mCherry-eGFP-W58X  -rrk1-crRNA-6-59 | mCherry-eGFP-W58X /*nmt1*  crRNA-pRNA(31bp)/*rrk1* | crRNA-eGFP-fusion-6-59-p5  crRNA-eGFP-fusion-6-59-p3 | 28+31 | 6 | This study |
| pDUAL-HFF-mCherry-eGFP-W58X  -rrk1-crRNA-6-65 | mCherry-eGFP-W58X /*nmt1*  crRNA-pRNA(37bp)/*rrk1* | crRNA-eGFP-fusion-6-65-p5  crRNA-eGFP-fusion-6-65-p3 | 28+37 | 6 | This study |
| pDUAL-HFF-mCherry-eGFP-W58X  -rrk1-crRNA-(-9)-65 | mCherry-eGFP-W58X /*nmt1*  crRNA-pRNA(A-9)/*rrk1* | crRNA-eGFP-fusion-(-9)-65-p5  crRNA-eGFP-fusion-(-9)-65-p3 | 28+37 | -9 | This study |
| pDUAL-HFF-mCherry-eGFP-W58X  -rrk1-crRNA-(-5)-65 | mCherry-eGFP-W58X /*nmt1*  crRNA-pRNA(A-5)/*rrk1* | crRNA-eGFP-fusion-(-5)-65-p5  crRNA-eGFP-fusion-(-5)-65-p3 | 28+37 | -9 | This study |
| pDUAL-HFF-mCherry-eGFP-W58X  -rrk1-crRNA-(-4)-65 | mCherry-eGFP-W58X /*nmt1*  crRNA-pRNA(A-4)/*rrk1* | crRNA-eGFP-fusion-(-4)-65-p5  crRNA-eGFP-fusion-(-4)-65-p3 | 28+37 | -9 | This study |
| pDUAL-HFF-mCherry-eGFP-W58X  -rrk1-crRNA-2-65 | mCherry-eGFP-W58X /*nmt1*  crRNA-pRNA(A+2)/*rrk1* | crRNA-eGFP-fusion-(2)-65-p5  crRNA-eGFP-fusion-(2)-65-p3 | 28+37 | 2 | This study |
| pDUAL-HFF-mCherry-eGFP-W58X  -rrk1-crRNA-12-65 | mCherry-eGFP-W58X /*nmt1*  crRNA-pRNA(A+12)/*rrk1* | crRNA-eGFP-fusion-(12)-65-p5  crRNA-eGFP-fusion-(12)-65-p3 | 28+37 | 12 | This study |
| pDUAL-HFF- mCherry-eGFP-W58X  -rrk1-crRNA-15-65 | mCherry-eGFP-W58X /*nmt1*  crRNA-pRNA(A+15)/*rrk1* | crRNA-eGFP-fusion-(15)-65-p5  crRNA-eGFP-fusion-(15)-65-p3 | 28+37 | 15 | This study |
| pDUAL-HFF- mCherry-eGFP-W58X  -rrk1-crRNA-19-65 | mCherry-eGFP-W58X /*nmt1*  crRNA-pRNA(A+19)/*rrk1* | crRNA-eGFP-fusion-(15)-65-p5  crRNA-eGFP-fusion-(15)-65-p3 | 28+37 | 19 | This study |

**TABLE S6.** crRNA-pRNA constructs for editing of endogenous genes.

| Plasmid Name | crRNA-pRNA primers | Promoter | Original plasmid | Targeting gene | Source |
| --- | --- | --- | --- | --- | --- |
| pDUAL-HFF-mCherry-eGFP-W58X  -rrk1-crRNA-act1-1566 | crRNA-act1-1566-p5  crRNA-act1-1566-p3 | *rrk1* | pDUAL-HFF-mCherry-eGFP-W58X | *act1* | This study |
| pDUAL-HFF-mCherry-eGFP-W58X  -rrk1-crRNA-ade6-622 | crRNA-ade6-622-p5  crRNA-ade6-622-p3 | *rrk1* | pDUAL-HFF-mCherry-eGFP-W58X | *ade6* | This study |
| pDUAL-HFF-mCherry-eGFP-W58X  -rrk1-crRNA-ade6-1003 | crRNA-ade6-1003-p5  crRNA-ade6-1003-p3 | *rrk1* | pDUAL-HFF-mCherry-eGFP-W58X | *ade6* | This study |
| pDUAL-HFF-mCherry-eGFP-W58X  -rrk1-crRNA-ade6-1160 | crRNA-ade6-1160-p5  crRNA-ade6-1160-p3 | *rrk1* | pDUAL-HFF-mCherry-eGFP-W58X | *ade6* | This study |
| pDUAL-HFF-mCherry-eGFP-W58X  -rrk1-crRNA-erp5-672 | cRNA-erp5-672-p5  cRNA-erp5-672-p3 | *rrk1* | pDUAL-HFF-mCherry-eGFP-W58X | *erp5* | This study |
| pDUAL-HFF-mCherry-eGFP-W58X  -rrk1-crRNA-mel1-921 | crRNA-mel1-921-p5  crRNA-mel1-921-p3 | *rrk1* | pDUAL-HFF-mCherry-eGFP-W58X | *mel1* | This study |
| pDUAL-HFF-mCherry-eGFP-W58X  -rrk1-crRNA-mug45 | cRNA-mug45-529-p5  cRNA-mug45-529-p3 | *rrk1* | pDUAL-HFF-mCherry-eGFP-W58X | *mug45* | This study |
| pDUAL-HFF-mCherry-eGFP-W58X  -rrk1-crRNA-nmt1-648 | cRNA-nmt1-648-p5  cRNA-nmt1-648-p3 | *rrk1* | pDUAL-HFF-mCherry-eGFP-W58X | *nmt1* | This study |
| pDUAL-HFF-mCherry-eGFP-W58X  -rrk1-crRNA-tdh1-79 | crRNA-tdh1-79-p5  crRNA-tdh1-79-p3 | *rrk1* | pDUAL-HFF-mCherry-eGFP-W58X | *tdh1* | This study |
